# An Epitope-Focused Trypanosome-Derived Vaccine Platform Elicits High-Affinity Antibodies and Immunity Against Fentanyl Effects

**DOI:** 10.1101/2022.05.02.487070

**Authors:** Gianna Triller, Evi P. Vlachou, Hamidreza Hashemi, Monique van Straaten, Johan Zeelen, Yosip Kelemen, Carly Baehr, Cheryl L. Marker, Sandra Ruf, Anna Svirina, Sebastian Kruse, Andreas Baumann, Aubry K. Miller, Marc Bartel, Marco Pravetoni, C. Erec Stebbins, F. Nina Papavasiliou, Joseph P. Verdi

**Author notes:** Dr. Joseph P. Verdi ( or). Lead Contact: Further information and requests for resources and reagents should be directed to and will be fulfilled by the lead contact.

## Abstract

Poorly antigenic small molecules pose challenges for the production of clinically efficacious antibodies. To address this, we have developed an immunization platform derived from the antigenic surface coat of the African trypanosome. Through sortase-based conjugation of antigens to the trypanosome surface coat protein variant surface glycoprotein (VSG), we created the VAST (**V**SG-immunogen **A**rray by **S**ortase **T**agging). We used VAST to elicit protective immunity to the effects fentanyl. Immunizing mice with fentanyl-VAST generated serological memory and protection from fentanyl challenge. Employing single-cell RNAseq, we then developed a streamlined method that synergizes with the VAST to identify immunization-elicited memory B cells and the antibodies they encode. All antibodies selected by this method displayed picomolar affinities for fentanyl. Passive immunization protected animals from fentanyl, while crystallography revealed that the mAbs bind fentanyl in an unusually deep pocket suited to small molecules, demonstrating the ability of the VAST to elicit high-quality therapeutic antibodies.

## Introduction

The human immune system struggles to generate protective antibody responses against small molecules unless they are fused to a larger immunogenic carrier protein (e.g., KLH, tetanus toxoid, etc.) to facilitate CD4+ T cell-dependent B cell activation, and combined with adjuvants prior to injection to prompt antigen recognition and increase immunogenicity. Classical hapten-carrier systems have been extensively tested in animals (Baruffaldi *et al*., 2019; Sen-Kilic *et al*., 2019; Deng *et al*., 2021). More recently, the desire to elicit antibodies to small molecule drugs of abuse has propelled implementation of classical hapten-carrier systems to humans with very limited success, and some prominent failures mostly due to the high individual variability in polyclonal antibody responses (Hoogsteder *et al*., 2012; Kosten *et al*., 2014). However clinical trials of nicotine and cocaine vaccines provided clinical proof of efficacy in those immunized subjects that met optimal levels of drug-specific antibodies.

Here, we describe a new antibody elicitation platform that avoids the need for exogenous adjuvants. This platform is extraordinarily versatile in that it can be conjugated with ease to virtually any molecule class of interest and reproducibly elicits strong antibody responses in mice. The system exploits the inherent immunogenicity of the blood-resident parasitic protozoan *Trypanosoma brucei* that largely depends on the densely patterned array of Variant Surface Glycoprotein (VSG) molecules that coat the surface of the organism (∼10 million copies of a specific VSG per trypanosome, forming the overwhelming majority of the cell’s total surface protein (Shimogawa et al., 2015)). This repetitive array facilitates epitope presentation to the immune system, driving both T-dependent and T-independent protective antibody responses (Reinitz and Mansfield, 1990; Verdi *et al*., 2020). Furthermore, antibody responses to VSG arrays are remarkably restricted in variable gene usage through an “epitope-focusing” effect that is potentially unique to this organism (see companion paper (Gkeka *et al*., 2021)).

In addition to epitope-focusing, VSG arrays also elicit long lasting responses: antibodies raised to a clonal VSG array protect the animal from infection with the same VSG-coated parasite for the life of the animal (Mugnier, Cross and Papavasiliou, 2015). Counterintuitively, long-standing evolutionary pressures have likely selected for VSG coats that elicit strong but highly VSG-specific antibody responses. This is due to the remarkable system of antigenic variation utilized by this pathogen. Trypanosomes possess a large genomic cache of antigenically distinct VSGs. During infection, the pathogen produces successive populations of “switched” cells that express different VSGs on the surface. VSG proteins with unique and immunodominant epitopes generate antibody repertoires that are highly focused on a given VSG (see companion paper (Gkeka *et al*., 2021)), but are unlikely to recognize switched cells. The host immune system thus invests considerable resources into a given VSG, efficiently clears parasites expressing it, only to find itself naïve to switched cells with different VSGs. Therefore, the VSG array that defines the coat structure of the parasite has been selected precisely for the ability to generate robust serological responses.

Over a decade ago, we explored whether we could derivatize the VSG-arrays that comprise the surface-coat of *T. brucei* with haptens of interest, and exploit their exceptionally antigenic nature. To assess if this was feasible, we genetically engineered VSGs to incorporate short peptides (e.g., FLAG) into their solvent-exposed loops with substantial success (Stavropoulos and Papavasiliou, 2010). To broaden the applicability of the system to other types of antigens, we then engineered VSGs to become substrates of the transpeptidation reaction known as “sortagging”. The term refers to the activity of enzymes produced by Gram-positive bacteria called sortases (Susmitha, Bajaj and Madhavan Nampoothiri, 2021). By genetically engineering sortaggable VSGs (Pinger, Chowdhury and Papavasiliou, 2017) and chemically synthesizing sortaggable antigens of interest (Figure 1A) we effectively converted the surface envelope of the trypanosome into a molecular display platform (Figure 1B). We can thus exhibit a wide variety of antigens against which antibodies are to be elicited (Plug-and-Display Platform, Figure 1C). We have named this platform the “VAST” (**V**SG-immunogen **A**rray by **S**ortase **T**agging) system. As a proof-of-concept, we sought to apply it to a clinically relevant target space that is both in need of alternative therapeutic options, and represents an antigen class that is populated by poorly antigenic target molecules.

**Figure 1.**
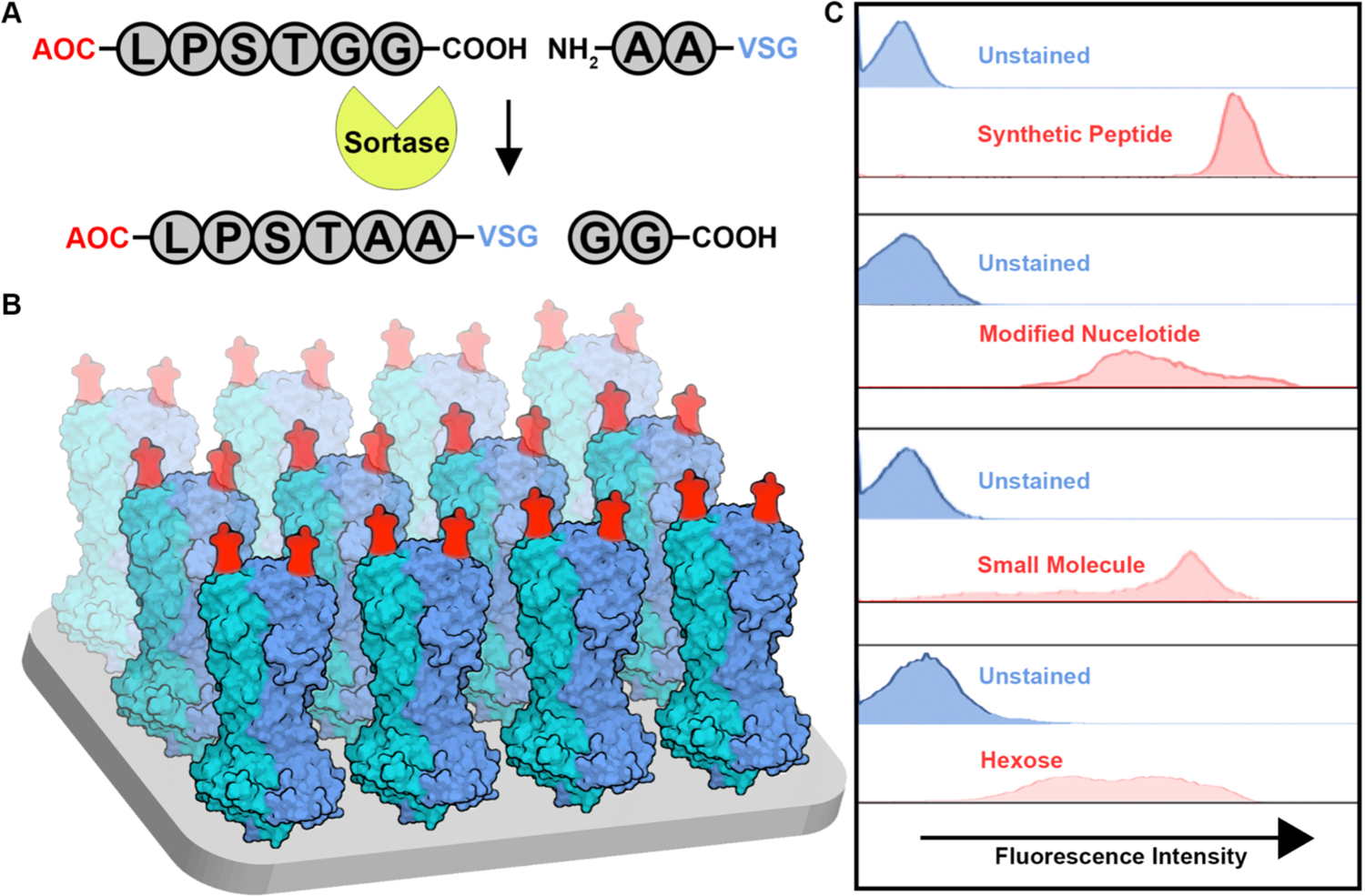
The VAST (VSG-immunogen Array by Sortase Tagging) platform: a trypanosome-based antigen display platform. **A.** Schematic of the VAST-specific sortagging reaction. Antigens of choice (AOCs) must be synthesized as a conjugate to the sortase recognition motif LPXTGG (X=S in this case), where the terminal glycine must have a free C-terminus. Sortase enzyme conjugates these AOCs to modified VSG proteins that contain an N-terminally extended AA motif. **B.** Model of the VAST platform highlighting the uniformity and density of the antigen-coated surface, making it a carpeted array for antigen presentation. Red moieties represent the sortagged antigens. The blue and teal structures are the VSG homodimers (each monomer represented by one color). Each VSG monomer has one N terminus and thus one sortaggable site, resulting in each dimer carrying a maximum of two copies of the AOC. C. Flow cytometry plots revealing the wide range of molecular classes that can be coupled to the VAST platform. Depicted here are the results from sortagging experiments using live trypanosomes sortagged with antigens representing the indicated molecule classes and detected using a variety of reagents (see methods for additional details).

Herein we have applied the VAST platform toward the elicitation of antibodies and immunity to the small molecule drug fentanyl. Fentanyl is a synthetic opioid 50-100-fold more potent than morphine. Although used clinically as a pain reliever and anesthetic, fentanyl is often mixed with recreationally used heroin or cocaine to mask impurities. This means that individuals are often unaware that they are self-administering the more potent fentanyl (Seth *et al*., 2018). Fentanyl is currently a leading cause of overdose and is the strongest driver of the increasing opioid-associated death rate. Death by opioid overdose was listed as the sixth largest cause of mortality in the United States in 2017, with figures having risen substantially during the COVID-19 pandemic (CDC, 2021).

Immunotherapeutics for substance use disorders (SUD) and overdose have been in preclinical development for some time (e.g., cocaine, heroin and other morphine derivatives, oxycodone, and fentanyl); the current status of vaccine development in the context of SUD is reviewed here (Pravetoni and Comer, 2019). Their development reflects both the acute need for relapse protection measures (the rate of relapse is > 80% after exiting rehabilitation clinics (Smyth et al., 2010; Zanda, Floris and Sillivan, 2021)), and the need for an adjunct to treatment with naloxone, the opioid receptor antagonist which is the current standard of care for reversal of overdose. Unfortunately, the short half-life of naloxone can lead to individuals “re-overdosing” due to “re-narcotization” from the same initial dose of fentanyl (Seth *et al*., 2018). To date, several vaccine formulations have shown efficacy against the pharmacological effects of fentanyl and its analogs including self-administration, antinociception, respiratory depression, bradycardia, and lethality (Torten *et al*., 1975; Bremer *et al*., 2016; Hwang *et al*., 2018; Raleigh *et al*., 2019; Robinson *et al*., 2020; Crouse *et al*., 2022). However, all of these efforts have employed the classical adjuvanted hapten-carrier protein combinations that have yet to produce a clinically approved product. These strategies have employed monovalent prime-boost combinations (i.e., using the same hapten-carrier protein conjugate for each injection), bivalent combinations employing the co-administration of admixed individual conjugates (Barrientos *et al*., 2021), and bivalent heterologous prime-boost combinations (Crouse *et al*., 2022) to target individual or multiple opioid compounds simultaneously. These strategies have also been used to generate mAbs against a variety of target drugs of abuse including fentanyl. Notably, fentanyl-specific mAbs are effective in both preventing and reversing fentanyl’s pharmacological effects in rodent models, further highlighting the promise of immunotherapeutics in the opioid space (Smith *et al*., 2019; Baehr *et al*., 2020, 2022; Ban *et al*., 2021). While promising indeed, there remains a lack of clinical output in this space and a desperate need for additional intervention options. Here, we have developed the Fent-VAST as a new tool, which consists of an extraordinarily large membranous hapten carrier that differs substantially in composition from conventional protein carriers (e.g., KLH). We have demonstrated its efficacy as an adjuvant-free vaccine against fentanyl. We then characterized the B cell response elicited by the platform and generated a series of high-affinity fentanyl-binding monoclonal antibodies (mAb) that could serve as overdose therapeutics in the clinic.

## Results

### The VAST (VSG-immunogen Array by Sortase Tagging) platform: a trypanosome-based antigen display platform

At present it remains a challenge to reliably induce protective antibodies using certain epitopes or antigens, especially when the antigen is a small molecule. These poorly immunogenic molecules are often coupled to larger antigenic carriers in order to elicit the desired responses. Here, we have developed a novel carrier called the VAST platform. VAST exploits the densely carpeted trypanosome surface coat through controlled attachment of antigens to the VSG proteins for antibody generation. To create the VAST platform, we took advantage of two pathogen-derived tools: (1) the VSG-array that covers the surface of each *T. brucei* cell, and (2) an enzyme that ligates peptidoglycan in the cell wall of Gram+ bacteria, the sortase transpeptidase. In principle, sortase will ligate polypeptide sequences as long as they each carry appropriate recognition motifs. We have used recombinant sortase enzyme to ligate an antigen of choice (AOC) to genetically engineered sortaggable VSGs carrying an N-terminal double alanine or double glycine motif extended above the surface coat by a flexible linker. The AOCs have been derivatized with the sortase recognition motif, LPSTGG with a free C-terminal end (Figure 1A). Sortase A recognizes both AOC-LPSTGG and AA-VSG, cleaves between the T and GG of the former (AOC-LPST*), and catalyzes the formation of a covalent bond between the T and AA-VSG, thus attaching the AOC to the VSGs on the surface of the organism (Figures 1A and 1B). Due to the uniquely dense coverage of the trypanosome surface by VSGs, a dense pattern of the conjugated antigen is thereby presented on the surface (Figure 1B). Using this approach, the surface envelope of the African trypanosome was converted into a dense molecular display platform; the VAST. The VAST has been successfully conjugated to a series of different molecules, representing a wide range of potential antigens to which antibodies could eventually be elicited (Figure 1C).

### The VAST platform induces antibody titers protective against fentanyl effects

To assess the effectiveness of VAST using a clinically relevant small molecule antigen, we chose the small molecule fentanyl. A sortaggable version of fentanyl (fen-sort) was produced using a modified synthesis of a detoxified fentanyl derivative previously described in Raleigh et al. and Robinson et al. (fentanyl and the synthesis scheme of the modified derivatives relevant to this manuscript are shown in Suppl. Figures 1A and B) (Raleigh *et al*., 2019; Robinson *et al*., 2020; Crouse *et al*., 2022). After sortase A-mediated conjugation of fen-sort to the surface VSG (Fent-VAST), sortagging efficiency was assessed by flow cytometry using a mouse anti-fentanyl mAb (Baehr *et al*., 2020, 2022). This revealed that 100% of the trypanosomes had been decorated with the fent-sort (Figure 2A).

**Figure 2:**
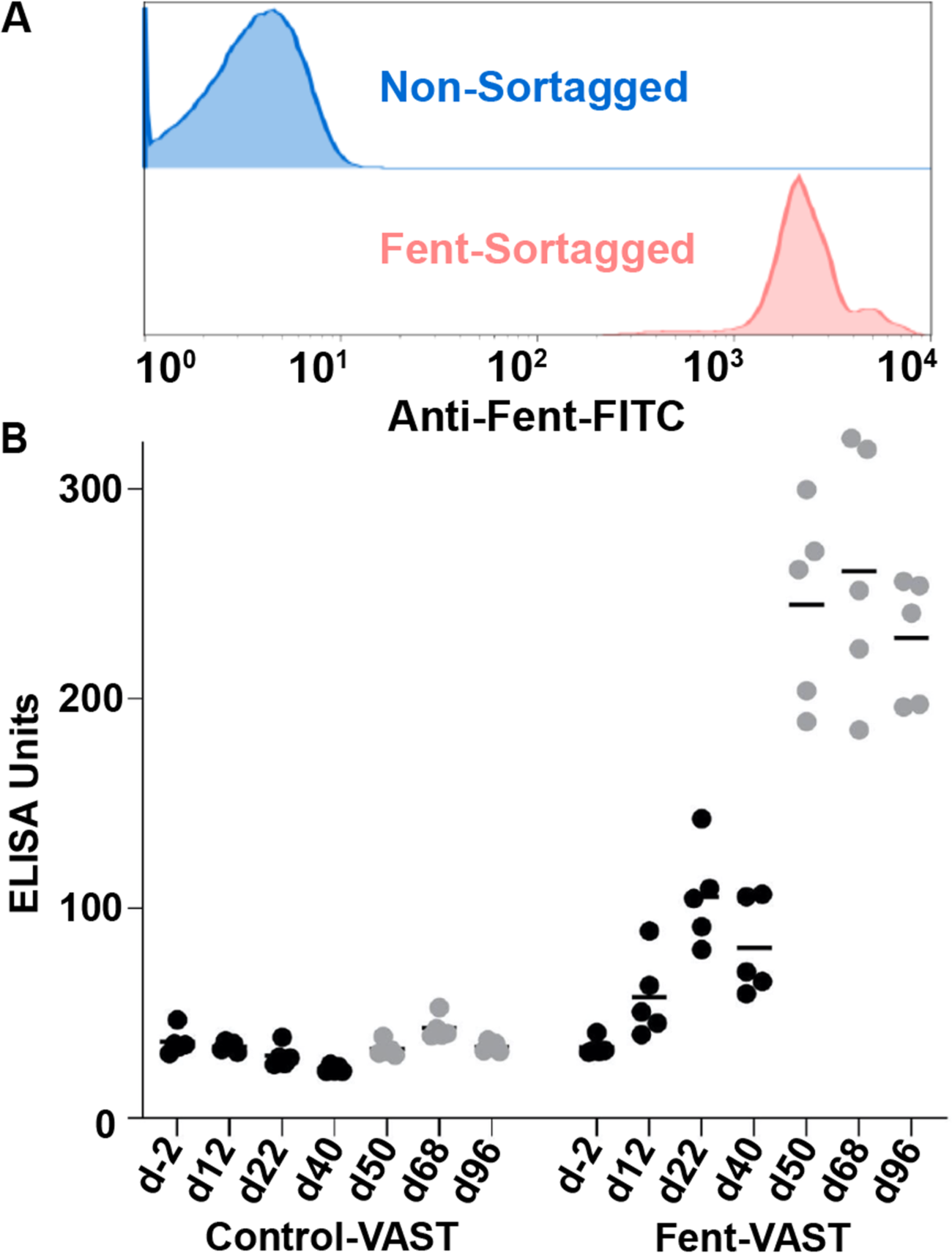
The VAST platform induces fentanyl-specific IgG antibody titers. **A.** After sortase A-mediated conjugation of the fentanyl-LPSTGG to VSG on the surface of trypanosomes, a mouse monoclonal antibody against fentanyl (provided by M. Pravetoni, University of Minnesota) was conjugated to FITC using a kit (Abcam, ab102884) and used to stain the sortagged trypanosomes followed by flow cytometry analysis using FACS-Calibur. Non-tagged trypanosomes were used as control for background staining. One representative plot shown. **B.** ELISA measurements of serum IgG against fentanyl are shown. The ELISA plates were coated with fentanyl-BSA at a concentration of 10 µg/ml. Sera were used at a starting dilution of 1:100 and diluted 2-fold. Plots show normalized AUC values (ELISA Units). The immunogen VAST or Fent-VAST is noted on the x-axis. Each dot represents antiserum from 1 mouse (n=5 mice per immunogen) (7 time points per immunogen starting with pre-bleed - collected at day-2 - up to post-2nd boost). The data shown in black circles were data collected prior to the first boost/recall injection, while data shown in grey circles were collected post-boost/recall. In this particular experiment, the first boost was delivered at day 42.

We have previously reported that immunizing mice with UV-crosslinked (inactivated) parasites recapitulated the immune responses launched by infection with the live organism (Pinger, Chowdhury and Papavasiliou, 2017). Thus, to assess the immune induction capacity of Fent-VAST, mice were immunized with either the Fent-VAST vaccine (comprising the material in Figure 1B in UV-inactivated form) or control-VAST (UV-inactivated but not sortagged).

Compelling evidence suggests that B cells normally encounter antigens in a membrane-bound format *in vivo* during initial activation (Carrasco and Batista, 2006), while we and others hypothesize that memory recall after initial activation can be quickly stimulated by soluble antigens since smaller entities are more easily able to penetrate B cell follicles (Roozendaal *et al*., 2009; Akkaya, Kwak and Pierce, 2020). Therefore, we designed a two-step injection paradigm where animals were “primed” with the membrane bound Fent-VAST array. Priming was followed by “boosting” with a 10- to 20-fold higher dose of antigen-conjugated VSG in a soluble format (i.e., after cleavage from the membrane and biochemical purification of the VSG protein; Suppl. Figure 2A-C). Blood was collected at several time points before and after the injections for antibody titer assessments. This prime/boost combination with two identical hapten-carrier entities displayed within distinct biophysical contexts (membrane bound array vs. soluble protein) led to the generation of high antibody titers against fentanyl in the mice immunized with Fent-VAST, but not the control-VAST, that matured with each injection (Figure 2B). Injection of membrane bound Fent-VAST alone led to the generation of only modest anti-fentanyl titers (Figure 2B, days 12-50), while injecting soluble-Fent-VAST in the absence of a priming step did not elicit high fentanyl titers (Suppl. Figure 2D). However, the combination of both steps resulted in the recalling of antigen-specific memory B cells that were produced after priming (Figure 2B), where recall is represented by the marked increase in titers (d50) after the first boost, which was administered on day 42. This ability to recall with soluble-Fent-VAST was maintained until at least 4 months post-priming (Suppl. Figure 2E; longer durations were not assessed), confirming that long-term immunological memory is established by the VAST platform. In all, this prime-boost combination was able to induce high fentanyl-specific antibody titers.

### Anti-fentanyl titers protect from fentanyl effects

Next, we tested whether the strong antibody response induced by the vaccine could protect mice from a behaviorally active dose of fentanyl. To this end, the antinociceptive activity of fentanyl was assessed using the hotplate assay as well as the Straub-Tail reaction test (assays described in Suppl. Figures 3A-C) 10 days after the final boost (vaccination schedule shown in Suppl. Figure 2C). Fent-VAST immunization reproducibly ablated any detectable fentanyl-induced antinociceptive effects via the hotplate test and fully prevented the typical Straub-Tail reaction (Figure 3A shows the results of one experiment involving 5 mice that is representative of multiple similar trials; Suppl. Figure 2C graphically illustrates the Straub-Tail response; videos show the Straub-Tail reaction of non-protected mice vs protected mice). This demonstrates that Fent-VAST vaccinated mice were protected from fentanyl toxicity whereas control mice were visibly behaviorally activated at the same dose (Figure 3A).

**Figure 3:**
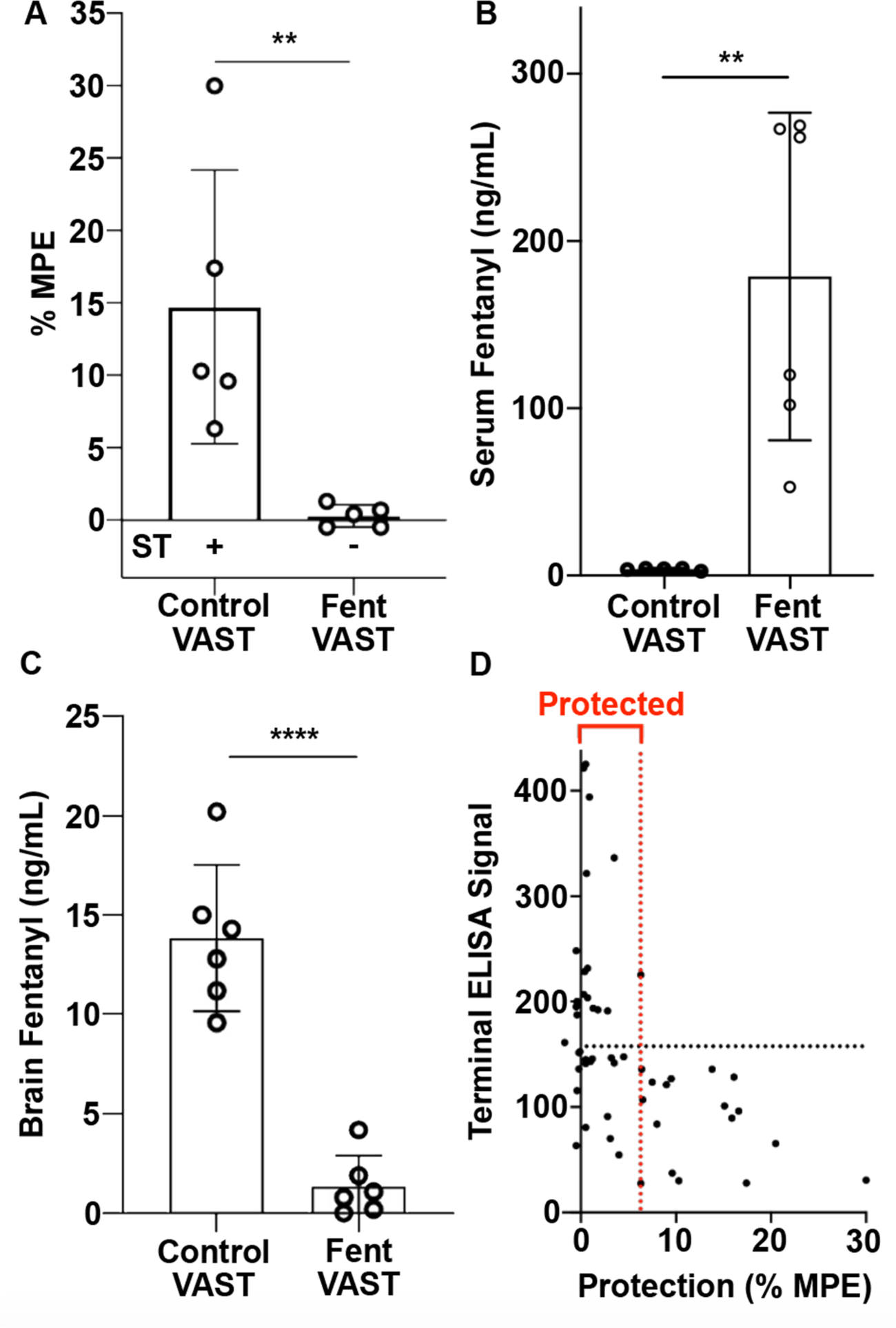
Anti-fentanyl titers protect from fentanyl pharmacological effects. **A.** Analgesic activity in mice immunized with carrier only (VAST) and in mice immunized with Fent-VAST at d100 was tested by using the hotplate antinociception assay as described by Cox and Weinstock (1964). The percentage maximum possible effect (%MPE) is shown. A positive (+) or negative (-) reaction in the Straub Tail (ST) test is indicated below and representative of the whole group. Means ± standard deviation of 5 mice per group are shown. (**) p < 0.01 by unpaired T test. **B-C.** GC-MS analysis of the distribution of fentanyl in the serum (B) and brain (C) is shown. Fentanyl concentrations were measured in mice immunized with control-VAST or Fent-VAST after performing a fentanyl challenge as described in Suppl. Figure 2A. Means ± standard deviation of 6 mice per group are shown. (****) p < 0.001, (**) p < 0.01 by unpaired T test. **D.** Correlation between anti-fentanyl serum titers and protection from fentanyl challenge in the antinociception assay is shown. Each dot represents an individual mouse. Mice shown here represent data collected from multiple similar vaccination trials. The comparison of terminal antibody titers (Y-axis) as determined by ELISA (normalized ELISA units) against the protection in behavioral experiments (X-axis, % MPE) suggests a protection cut off of approximately 175 ELISA units (black dashed line). Mice that did not display any fentanyl-induced antinociceptive effect (< 7%MPE - red dashed line) fall under the “protected” bracket. All mice with a normalized terminal ELISA titer >175 ELISA units fall into this category, while mice with terminal ELISA titers that do not meet this cutoff were not reproducibly protected from fentanyl intoxication.

In theory, antibodies are thought to protect against fentanyl effects by binding to the drug in the bloodstream resulting in decreased circulating unbound (free) drug. This is hypothesized to prevent fentanyl from entering a multitude of tissues (Crouse *et al*., 2022), although the most clinically relevant target space is the central nervous system. We thus measured the fentanyl concentration in the brain and serum of mice after fentanyl challenge by LC-MS. While control mice exhibited levels of approximately 10-18 ng/ml fentanyl in the brain and less than 5 ng/ml in the serum, the Fent-VAST vaccinated mice exhibited less than 5 ng/ml in the brain and approximately 80-280 ng/ml fentanyl in the serum (Figures 3B and C). This demonstrates that the Fent-VAST-induced antibodies were capable of trapping the fentanyl in the serum of vaccinated mice, preventing the drug’s blood-brain-barrier penetration and ultimately the neuropharmacological effects seen in the control mice (videos). Finally, we injected naloxone immediately after the behavioral assessments (Suppl. Figures 3A and B). Fentanyl-induced antinociception was reversed in all mice as expected (Suppl. Figure 3B).

As a pre-clinical assessment of whether the immune response to the vaccine could be used as a predictor of efficacy, we assessed the degree to which fentanyl-induced antinociception was reduced in mice with a wide range of anti-fentanyl antibody titers. We were able to identify an ELISA titer value (175 ELISA units) that serves as a protection “cut-off” with this dose of fentanyl (100 µg/kg), whereby mice with titers higher than this value were protected while mice with a lower titer responded to fentanyl similarly to controls (Figure 3D). These data provide the conceptual basis for the clinical assessments that could eventually be established in order to predict vaccine efficacy in humans.

### The VAST platform elicits six different B cell subsets

Ideal vaccines not only provide short-term protection against the foreign entity, but also establish the formation of long-lasting immunological memory. Since the VAST platform could be used to produce vaccines as well as antibodies, we sought to assess whether the platform establishes *bona-fide* memory B cells, as would be predicted by the serology data (Figure 2B, Suppl. Figure 2E). To this end, we applied a transcriptomics-based approach using single cell RNA-seq to precisely identify the different B cell subsets induced by the VAST platform. Using flow cytometry, we sorted all fentanyl-binding B cells from two mice immunized with Fent-VAST (the gating strategy and baiting results are shown in Suppl. Figures 4A and B) and performed single-cell RNA sequencing. Based on the iNMF factor loadings (group of genes defining a biological signal) defined by integrative non-negative matrix factorization (iNMF) (Welch *et al*., 2019) and by applying the Uniform Manifold Approximation and Projection (UMAP) algorithm, we were able to identify six distinct B cell subpopulations among the sorted fentanyl-specific B cells (Figure 4A). The subpopulations were annotated based on the expression levels of the top 10 most differentially expressed genetic markers for each of the sub-cluster (Figure 4B), among them: Myc, Cd55, Cd21 (or Cr2), Cd80, and Fcrl5 (Riedel *et al*., 2020). To strengthen our confidence in the subcluster identification, we also examined the expression levels of 35 of the most widely recognized B cell markers such as Fcer2a (or Cd23), Mzb1, Cd9, Cd38, Ccr4, Fos, Cd40, and several transcription factors such as Irf4, Irf8, Blimp1, Bach2, and Bcl6 (Figure 4B) (Shi *et al*., 2015; Meng and White, 2017; Welch *et al*., 2019; Riedel *et al*., 2020; King *et al*., 2021; Laidlaw and Cyster, 2021). The subpopulations included a germinal center light-zone pre-plasma population (GC-LZ-pre-plasma), a dark-zone population (GC-DZ), a follicular B cell population (FOB), a marginal zone-like atypical memory B cell population (MZB), and a naïve B1b cell population (Cerutti, Cols and Puga, 2013; Pérez-Mazliah *et al*., 2018; Gonzales *et al*., 2021; Mathew *et al*., 2021). Importantly, there was also a recognizable isotype-switched memory B cell population (switched-MBC) identified with this analysis. These data further support earlier pre-clinical evidence that opioid-carrier conjugate vaccines induce CD4+ T cell-dependent B cell processes to elicit opioid-specific IgG antibodies, including germinal center formation as assessed by antigen-specific GC B cells, Tfh, GC-Tfh, and subsequent switched memory B cells (Laudenbach *et al*., 2015; Baruffaldi *et al*., 2018; Robinson *et al*., 2019; Crouse *et al*., 2020). Pairing this population with data from FACS index sorting, we were able to show that the cells with the highest affinity to fentanyl were contained in this isotype-switched memory B cell subpopulation (Figure 4A, right panel), which was also enriched for Cd80 and Cd73, two common T-dependent memory B cell markers that are absent from the other subpopulations (Tomayko *et al*., 2010). The Mki67 proliferation marker was also upregulated in the switched-MBC population, suggesting the reactivation of MBCs immediately after the boost administered 10 days prior to cell collection, which is consistent with the known kinetics of the recall response (Figure 4C and Suppl. Figure 4C) (Palm and Henry, 2019). Additionally, the subpopulation expresses longevity markers such as Zbdb20, traditionally expressed in long-lived plasma cells (Chevrier *et al*., 2014; Wang and Bhattacharya, 2014). The unswitched populations produced B cell receptors (BCRs) mainly of the IgM isotype, while the switched memory population predominantly expressed IgG1 and IgG2b (Figure 4D). To further characterize these BCRs from the switched-MBCs, we performed BCR repertoire analysis based on VDJ usage of the heavy and light chains of the BCRs. We found that many of the cells from the switched-MBC subpopulation showed a very similar VDJ profile for paired heavy and light chains (Figure 4E). In addition, they showed several silent and missense somatic hypermutations (SHM), another line of evidence suggesting that these cells underwent germinal center reactions and are thus memory B cells. Heavy and light chain pairing of the memory B cells was dominated by the VH1-74/JH4 or VH1-53/JH2 genes in combination with VK6-15/JK2 (Figure 4E), pairs that are not at all represented within the cumulative repertoire of the five non-memory subsets (Suppl. Figure 4D). Notably, these pairs are also not represented in naïve mouse BCR repertoires or repertoires from mice infected with live *T. brucei* (see companion paper (Gkeka et al., 2021)). In addition, these BCRs had very similar or even identical heavy and light chain CDR3 regions and shared several SHMs, suggesting that these cells are clonally related (Table 1). These analyses make clear that the vaccine platform was able to induce an isotype-switched fentanyl-specific memory B cell population that included cells with highly similar or even identical BCRs in separate mice.

**Figure 4:**
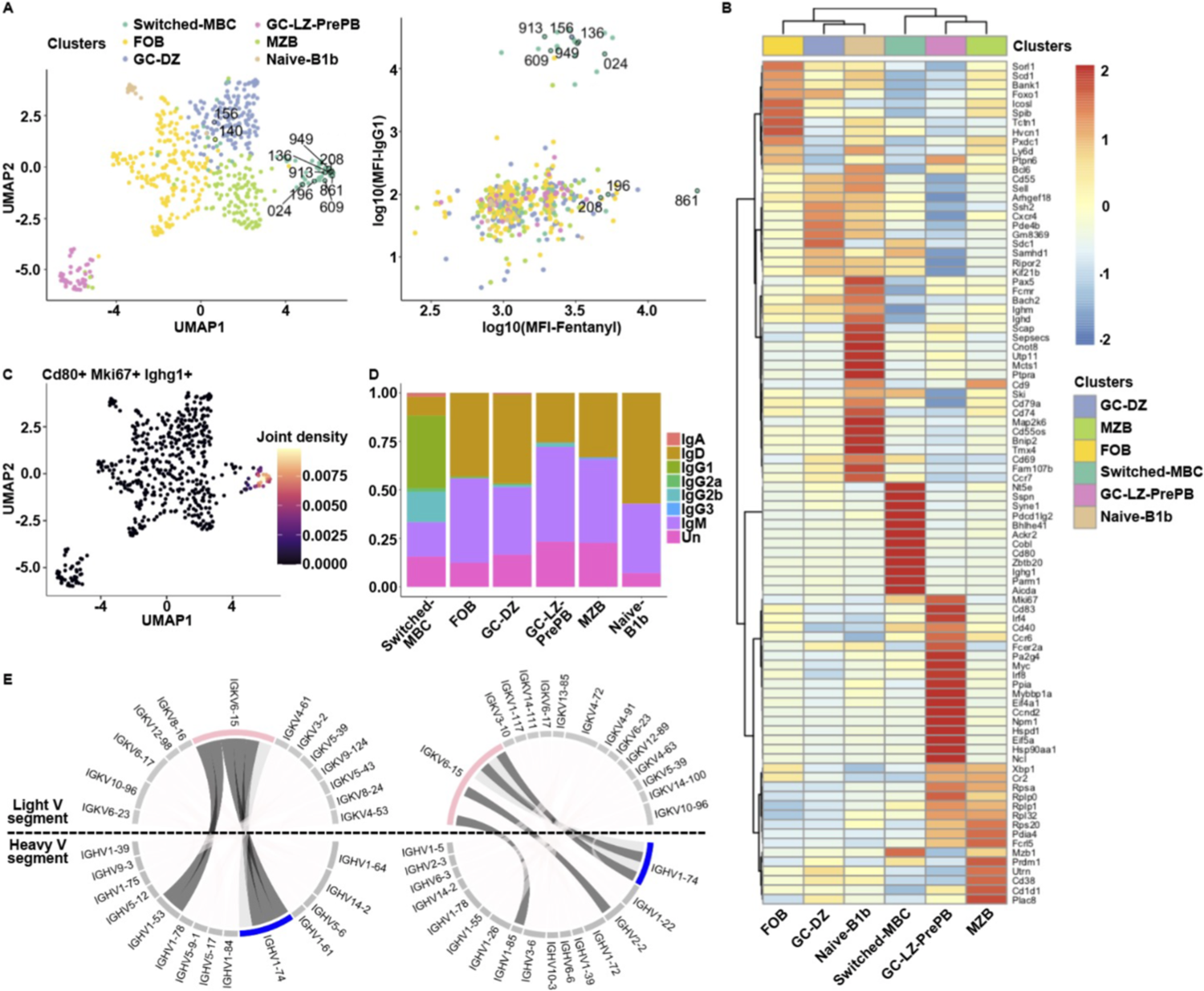
The VAST platform elicits six different B cell subsets. **A.** UMAP visualization of the B cell subclusters and scatter plot of the log10(MFI) for the fentanyl binding versus the IgG1 surface expression is shown. UMAP visualization of the iNMF analysis-resulting factors revealed that the Fentanyl specific B cell population is clustered into 6 distinct subpopulations based on the transcriptional expression. The color code indicates the different subclusters. The determined identities of the subpopulations are defined in the legend above the graph. The numerically-labeled cells correspond to the B cells from which the BCRs were cloned to produce Fabs and full-length antibodies. **B.** Heatmap visualizes the top 10 statistically significant upregulated genes based on the MAST statistical framework for each subcluster along with several known B cell markers. Genes have been identified as markers with an adjusted p-value < 0.05. The columns correspond to the different subclusters (the same color code used in 3A) and the rows to the average gene expression for the selected genes. Hierarchical clustering is applied in both rows and columns using the euclidean and correlation distance, respectively. Red stands for high gene expression and blue for low. **C.** UMAP visualizes the joint expression of Cd80, Mki67 and Ighg1 genes. The joint expression is calculated by multiplying the kernel density estimates from all 3 genes. Purple stands for low simultaneous expression while beige stands for high. **D.** Barplot depicts the heavy chain isotype distribution for each B cell subpopulation. There are cells (labeled as Unknown-Un) for which we were unable to identify the isotype via BLAST. Each color corresponds to an isotype, and each column to a B cell subcluster. **E.** Circos plots of the switched memory B cell subpopulation for each of the two mice are shown. The expanded heavy (in blue - IGHV1-74) and light (in pink - IGKV6-15) variable chain genes are highlighted. In dark gray we show the cells that were cloned based on their heavy and light chain gene combinations. In light grey we highlight the non-cloned BCRs from this paired group.

**Table 1:**
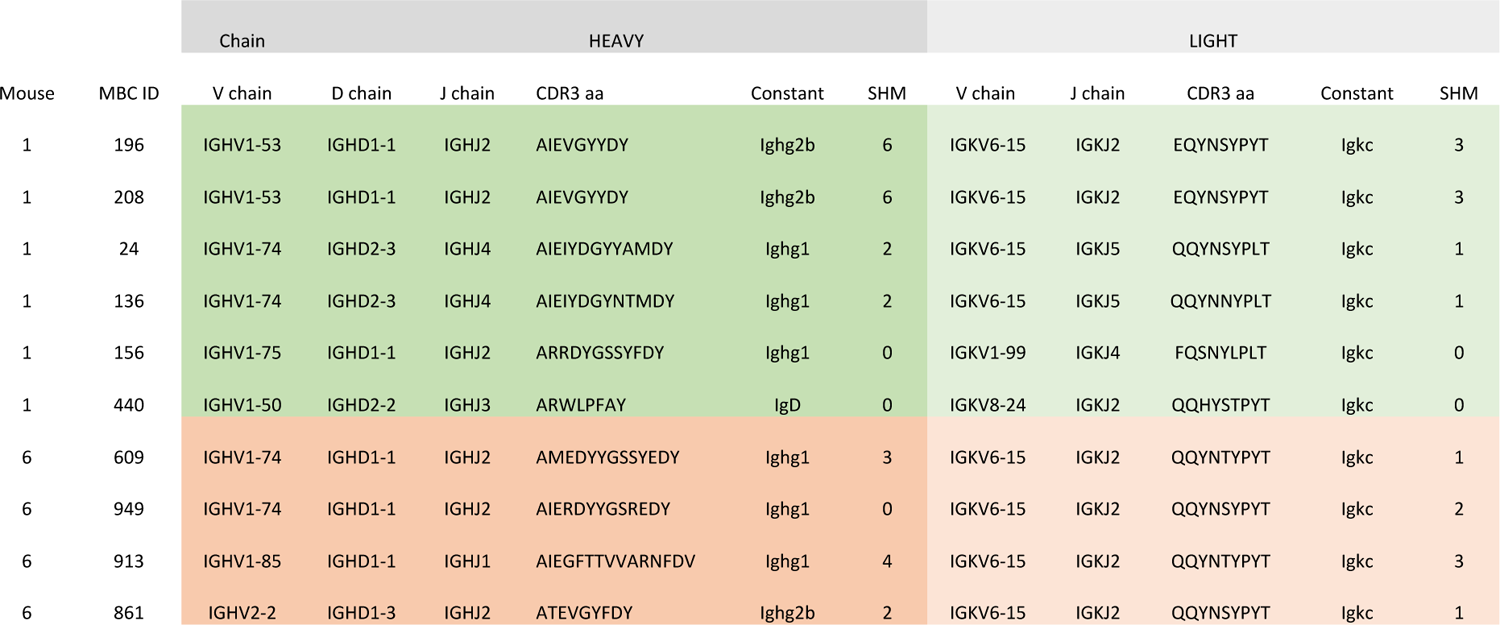
Monoclonal anti-fentanyl antibodies cloned from BCRs of B cells (MBC ID) belonging to the switched memory B cell population. The gene usage of the V(D)J combination is indicated, along with the CDR3 amino acid sequence, and the constant region for each heavy and light chain combination. In the “SHM” column the number of silent and missense mutations detected along the full length of the heavy or light chain is indicated.

### Fentanyl-specific antibodies are of high affinity, specificity, and protect mice from fentanyl effects

To assess the functional relevance of the antibodies encoded by these specific Ig genes from the isotype-switched MBC subpopulation (Table 1), we cloned and expressed some of the BCRs as recombinant full-length mAbs or Fabs and purified them (Figures 5A and B). Affinity and thermodynamic characteristics of purified Fabs binding to fentanyl-citrate and a fen-sort were first analyzed using isothermal titration calorimetry (ITC). All fentanyl binding reactions were exothermic with nanomolar and subnanomolar ranges of affinities (Figure 5C). Large enthalpy change (dH ∼ 68-77 kJ/ml, Fab208 - 100kJ/mol) coupled with unfavorable entropy input (13-23 kJ/mol, Fab208 - 47kJ/mol) contributed to a binding energy of approximately 50-60 kJ/mol. The binding reaction of fen-sort was similarly enthalpy driven (dH ∼ 132-156 kJ/mol, dS∼ 80-105 kJ/mol) and resulted in 0.7-1nM K_d_s. The subnanomolar affinities for both unmodified fentanyl-citrate and fen-sort fell below the optimal measurement range for ITC, and indeed the raw data suggested that the actual affinities may be higher than what had been calculated. We therefore used biolayer interferometry, a technique with a wider dynamic range, to measure the affinities of anti-fentanyl antibodies to the fen-sort. Similarly to ITC, these affinities were in the picomolar range, spanning from 980 pm to less than the measurable limit of the experimental apparatus (experimental limitations discussed in the methods section; Figure 5C). Competition ELISA confirmed that these antibodies were fentanyl-specific and importantly did not interact with naloxone (Figure 5D), suggesting that the Fent-VAST does not elicit antibodies that would interfere with naloxone if used in combination.

**Figure 5:**
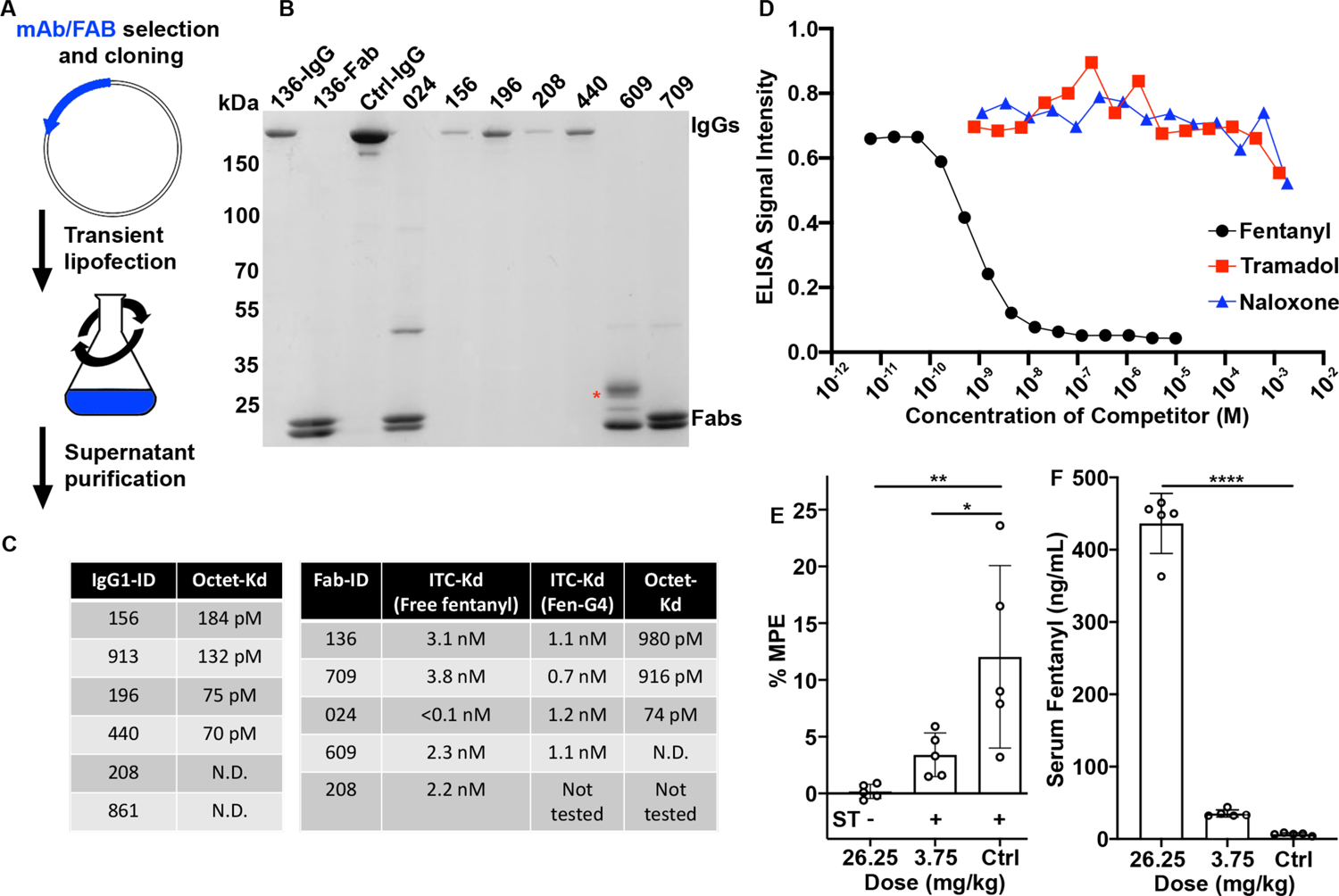
Fentanyl-specific antibodies are of high affinity, specificity and protect mice from fentanyl pharmacological effects. **A.** Schematic of the expression system used to produce recombinant antibodies. Plasmids encoding secretable heavy and light chain full length or Fab sequences are transiently expressed in suspended HEK cells prior to chromatographic purification from the supernatants. **B.** Coomassie-stained non-reducing SDS-PAGE after affinity purification of a panel of the expressed IgGs (running near 150 kDa) and Fabs (running as separate heavy and light chains near 25 kDa). The red asterisk marks a band produced by a glycosylated Fab, while the remaining Fabs are not glycosylated. The first lane marked “136” is the full-length IgG version of this antibody, while the second lane is the Fab. 156, 196, 208, and 440 are IgG protein samples, while 024, 609, and 709 are Fab protein samples. Ctrl-IgG is a control IgG sample as an additional molecular weight reference. **C.** Binding affinities determined for each antibody using biolayer interferometry with a biotinylated fentanyl hapten as the target molecule. The instrument can is limited in its ability to accurately determine an affinity in certain conditions; for example if the off-rate is too long. Antibodies that fall into this category are likely to also have high affinities (sub-nanomolar) but are here reported as “not determinable” (N.D.). **D.** Competition ELISA data generated with haptenated fentanyl-coated plates. The indicated soluble competitors are serially diluted into the assay to determine the relative efficiency of cross-binding to other opioid molecules, with soluble fentanyl serving as a positive control. **E.** Five 6-8 weeks old female C57BL/6J mice per group were passively immunized by intraperitoneal injection of 2 concentrations monoclonal antibody 136. One day later, mice were challenged in antinociception assays for their analgesic reactivity using the hotplate antinociception assay as described in Suppl. Figure 3. The percentage maximum possible effect (%MPE) and Straub Tail reaction are shown. A positive (+) or negative (-) reaction in the Straub Tail (ST) test is indicated below and representative of the whole group. Mean ± standard deviation of 5 mice per group are shown. (*) p < 0.05, (**) p < 0.01 by Dunnett’s multiple comparisons test. **F.** Serum from the mice in (E) were collected and analyzed for fentanyl content by mass spectrometry. (****) p < 0.001 by Dunnett’s multiple comparisons test.

To investigate if the memory B cells contributed to protection, we selected, cloned and expressed as soluble antibody FenAb136 (with a calculated affinity of 980 pM), derived from the BCR sequencing information of a memory B cell, to determine if the antibody was able to directly protect mice from fentanyl challenge. Mice were passively immunized by intraperitoneal injection with two different concentrations of the full-length antibody. One day later, immunized mice were challenged against fentanyl using the hotplate antinociception assay for their analgesic reactivity to fentanyl using the hotplate antinociception assay as described in Figure 3 and Suppl. Figure 3. Although only 37% (higher dose) and 25% (lower dose) of the originally injected amount of antibody could be detected in the blood by western blot at the time of fentanyl challenge (24h post immunization) (Suppl. Figure 5A), the mice were protected from fentanyl effects in a dose-dependent manner (Figure 5E). Mass spectrometric analysis (LC-MS/MS) showed that fentanyl was trapped in the serum of the mice administered with the antibodies as expected (Figure 5F). In fact, fentanyl could be detected at the same molar concentration in the serum as the antibody (approximately 1.2 µM for high dose and 0.1 µM for low dose; Suppl. Figure 4B). In summary, we were able to show that VAST-elicited antibodies are specific to fentanyl and are able to protect mice from a fentanyl challenge by trapping the fentanyl in the blood and preventing it from reaching its receptors in the brain. Therefore, these mAb could be used as another addition to the overdose-prevention toolkit as prophylactics to prevent overdose up to equimolar concentrations of drug.

### VAST-elicited antibodies bind fentanyl in a deep, enveloping pocket

In order to further characterize the antibodies produced by the VAST platform, we co-purified drug-Fab complexes (two with fentanyl and two with fen-G4) and determined four high-resolution structures by X-ray crystallography (Figure 6, Suppl. Figure 6, and Suppl. Table 1). All the antibodies share a similar binding mode, consisting of a deep, invaginated pocket formed by residues from both the heavy and the light chains of the immunoglobulin. While the upper portions of the pocket consist of CDR regions of the chains, the pocket is so deep (approximately 15 angstroms from the upper portions to the lower regions) that the bottom of the pocket is lined with residues from the beta-sheet framework regions, with several amino acids from these segments making contacts to the drug.

**Figure 6:**
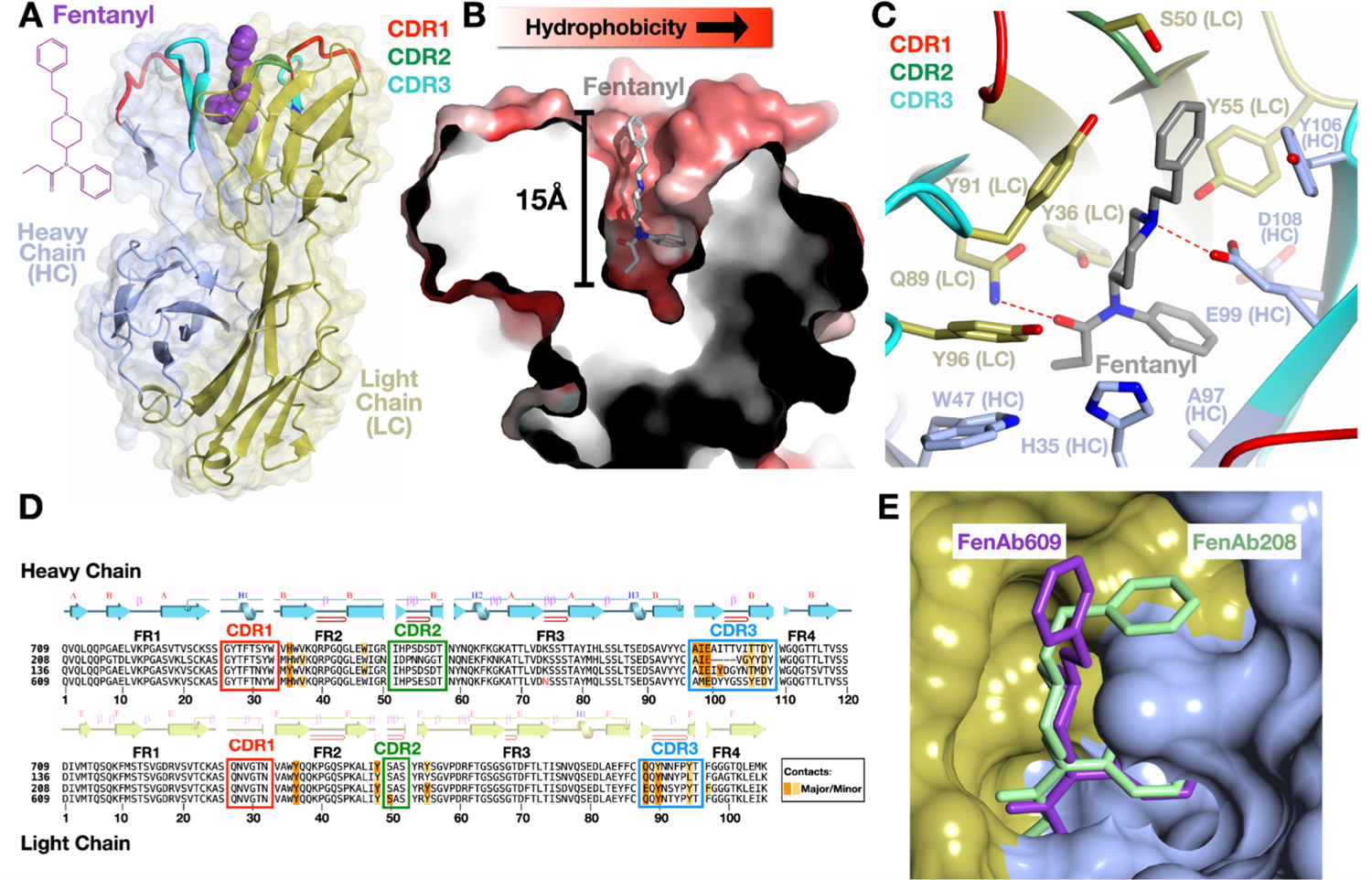
VAST-elicited antibodies bind fentanyl in a deep, enveloping pocket. **A.** Overall structure of a complex of a Fab (FenAb609, heavy chain colored light blue, light chain colored gold) with fentanyl (purple, space filling depiction) shown as a ribbon diagram with the two-dimensional chemical structure of fentanyl on the left. The CDR1, CDR2, and CDR3 regions of the antibody are colored in red, green, and cyan respectively. The partially transparent molecular surface of the protein is shown overlaid on the structure. **B.** Illustration of the fentanyl binding pocket as a thin slice through the molecular surface of the protein (colored in a gradient from white to red to reflect increasing hydrophobicity of the surface, colored using the method of Eisenberg (Eisenberg, D., Schwarz, E., Komaromy, M. & Wall, R. Analysis of membrane and surface protein sequences with the hydrophobic moment plot. J. Mol. Biol. 179, 125–142 (1984)). Fentanyl is shown as a stick model with atoms of carbon, nitrogen, and oxygens in gray, blue, and red, respectively. **C.** Contacts of the protein to fentanyl shown with side chains as stick models and the mainchain as a ribbon diagram. Fentanyl colored as in panel (B), and the residues of the heavy and light chains adopting the colors from panel (A). The CDR regions are colored as in panel (A) as well. “HC” denotes heavy chain and “LC” denotes the light chain. Hydrogen bonds are shown as dashed red lines between bonded atoms. **D.** Sequences of four fentanyl-binding Fab molecules where the alignment was generated by superimposing the crystal structures. The secondary structure of a representative Fab (FenAb609) is shown above the sequence, colored as per panel (A), as are CDR regions. Disulfide bonds are shown as lines connecting cysteine residues. Major and minor contacts are indicated in orange and yellow, respectively. The framework regions are denoted as FR1, 2, 3, and 4. **E.** Superposition of FenAb609 and FenAb208 crystal structures with the molecular surface of FenAb208 shown (heavy and light chain regions colored as panel (A)) along with the stick figures of fentanyl in both molecules to illustrate the different conformation of the top aromatic ring in the FenAb208 structure, which possesses a different CDR3 conformation which opens the molecular surface for the alternative conformation of the drug.

Fentanyl (N-(1-(2-phenylethyl)-4-piperidinyl)-N-phenyl-propanamide) inserts deeply into the cavity in a mostly elongated conformation (Figure 6, illustrating primarily the complex of FenAb609). Projecting only slightly out from the molecular surface of the Fab is the phenylethyl ring (the group absent in the fen-G4 construct used to elicit the immune response in mice; Suppl. Figure 1A vs 1B). This ring is mostly buried (Figure 6B), and the remaining chemical structure plunges straight into the pocket so that the piperidine ring is fitted tightly about halfway into the cavity, locked in place with numerous hydrophobic van der Waals contacts and a hydrogen bond (Figure 6C). The drug descends further into the pocket bottom where the two groups of the N-phenyl-propanamide group each plug into a snug series of contacts. A cavity is carved into the surface at the pocket bottom into which the benzene ring inserts, while the propionyl group projects in the opposite direction and is held tightly by numerous contacts including a deeply buried hydrogen bond. Several contacts from framework residues occur in this region at the bottom of the pocket (Figures 6C and D). The network of interactions observed in the structures support the ITC data, which suggested that the net formation of hydrogen bonds between fentanyl moiety and Fab was a major driving force of binding and antigen specificity. Interestingly, binding of ligand (e.g., testosterone, steroids, musk odorant) to a deep hydrophobic pocket on Fab is often hypothesized to be regulated by shape complementarity and the concomitant desolvation of the complementary surfaces with a subsequent rise in system entropy (Langedijk *et al*., 1999). Int he case of the fentanyl-specific antibodies described here, entropy change was unfavorable, indicating that conformational rearrangements of Fab loops were likely required to form the pocket around fentanyl. Unbound Fab should exhibit higher degrees of freedom which were lost upon ligand binding, leading to an entropy drop. This hypothesis correlates well with the observation from crystallization trials where the unliganded Fab could not be crystalized.

As noted, the pocket structure is common to all the antibodies identified, and while many contacts are conserved between the different Fabs, there are differences that could contribute to the range of affinities measured. In addition, the “top” aromatic group of fentanyl is seen in FenAb208 to adopt a different conformation than in some of the other fentanyl-bound antibodies (Figure 6E). In particular, the CDR3 loop of the heavy chain in FenAb208 is significantly shorter and adopts a very different conformation that creates a small socket not present in the other structures, into which the phenylethyl ring inserts. Because this ring is absent in fen-G4, it demonstrates that the antibody repertoire is able to produce a collection of different immunoglobulins that can engage this chemical group effectively, even without any pressure to select such variants.

## Discussion

Many obstacles remain to produce specific and high-affinity polyclonal antibodies and mAb against many different categories of antigens, one being small molecules. Here we have developed a trypanosome-based antigen presenting platform which can serve both as a vaccine and a mAb-producing tool, particularly in contexts where more conventional methods have failed. By sortagging the antigen of interest to the surface of trypanosomes, a method that retains the native architecture of the trypanosome surface coat, and injecting this construct into mice, the VAST platform can induce high antibody titers specifically against the sortagged antigen. The elicitation of these high titers does not require the addition of adjuvant, contrary to most conventional vaccination methods. Thus, the VAST serves as a “self-adjuvanting”, densely covered antigen presentation array that harnesses the natural immunogenicity of the trypanosome coat in order to elicit an immune reaction against even poorly immunogenic antigens. This is a significant advantage over conventional adjuvant-requiring vaccine platforms, as only few adjuvants are currently approved for human use (Petrovsky, 2015). Further, despite their approved status, many of these adjuvants are still known to cause adverse events and local side effects. While we have not yet formally assessed the toxicity of our formulations or performed pharmacokinetic studies, we expect that the lack of adjuvant incorporation will facilitate positive results in such studies. We thereby hypothesize that the VAST could have a reasonably straightforward route through clinical approval processes, since formulations will be kept standard with each new antigen linked to the platform. In addition, the design of the VAST platform makes it easily adaptable to not only different antigens but even different classes of antigen, as long as the antigen can be synthesized in a sortaggable format.

The ability to elicit antibodies against small molecules is dependent on a multitude of factors, including the stability and clearance half-life of a molecule of interest. Small molecules that are rapidly purged from the body may evade B cell detection for temporal reasons, while the molecules themselves are also unlikely to have the capacity to be presented on MHC or act as T cell epitopes. Free small molecules are thus unlikely to stimulate antibody responses. However, use of chronic opioids has been reported to elicit low levels of anti-opioid antibodies (Kyzer *et al*., 2020), but little is known about the underlying mechanisms permitting antibody elicitation. Thus, a core concept in the conventional design of small molecule vaccines is to conjugate the small molecule hapten to a larger immunogenic carrier that drives a CD4 T cell response in order to sustain B cell activation, expansion, and differentiation. In fact, depletion of CD4 T cells, use of TCR KO mice, and T cell independent carriers such as ficoll and dextran (Laudenbach *et al*., 2015; Baruffaldi *et al*., 2018) show that T cell help is required for the generation of antibody responses in the context of opioid vaccines. However, decades’ worth of literature show that trypanosomes drive a strong T independent antibody response (Reinitz and Mansfield, 1990; Verdi *et al*., 2020), and so we cannot rule out the roles that T independent pathways may have in the immune response to VAST-carried haptens at this time. Generally, we hypothesize that immune stimulation by the VAST is distinct from that which is driven by conventional protein carriers. For example, the VAST platform prime steps are, to our knowledge, the largest antigenic carrier currently under investigation. The inherent antigenicity of the trypanosome notwithstanding, the size of the organism is substantially larger compared to that of smaller carriers like KLH (an entire eukaryote versus a large protein). Classically, material of that size is likely to be cleared through phagocytosis, while we hypothesize that smaller carriers are unlikely to stimulate that type of clearance mechanism. The different mechanism by which antigens are recognized, cleared, and processed is likely to lead to noticeably different immune stimulating outcomes (e.g., different types of cytokine and chemokine responses). These differences may facilitate the strong responses driven by the VAST that we have observed in the absence of adjuvants.

In this study fentanyl is used as a proof-of-concept antigen as it represents a difficult antigen class (small molecules), has a critical medical application (protection against overdose), and possesses a mature collection of assays and behavioral studies. We demonstrate that Fent-VAST is able to induce high antibody titers against fentanyl that are protective against the behavioral effects of opioids. These antibodies appear to act as a “sponge” that soaks up the fentanyl molecules when they enter the bloodstream, thereby preventing them from reaching their target receptors in the brain. A protective antibody titer threshold was determined that could form the basis for a strategy to predict vaccination success in the clinic. Clinical assessments of nicotine and cocaine vaccines also identified titer thresholds required for protection, although unfortunately only a minor percentage of participants responded well enough to eclipse that threshold in clinical trials (Hoogsteder *et al*., 2012; Kosten *et al*., 2014), leading to the abandonment of these projects. It should be noted, however, that the major targeted outcome in these trials was cessation of drug use and thus the overcoming of addiction. This is in stark contrast to the concept of an overdose-preventing vaccine, which strives to prevent death rather than prevent substance use disorder entirely. A vaccine of this nature would thereby need to be combined with additional existing substance use disorder mitigation strategies, although we hypothesize that a clinical outcome of mortality prevention may be easier to achieve than tackling addiction itself, and thereby maintain that opioid vaccine research has an important role to play in the future of this field.

Importantly, we show that the VAST platform is able to induce memory B cells, which is a crucial quality checkpoint for vaccines. Antibody titers usually drop with time, but memory B cells can persist for long periods and be reactivated upon an additional encounter with the same antigen, differentiating into plasma cells and contributing to antibody titers. Previous studies have shown that conjugate vaccines elicit opioid-specific B cell population subsets including switched memory B cells (Laudenbach *et al*., 2015), and that B cell formation is dependent upon germinal center formation and involvement of cognate CD4+ T cells (Laudenbach *et al*., 2015; Baruffaldi *et al*., 2019). Furthermore, vaccination could boost antibodies well-beyond the disappearance of the first antibody response (Raleigh *et al*., 2021). Here, VAST-elicited memory B cells are persistent as they can be recalled in mice through at least 16 weeks after the second prime injection, which renders this platform flexible for different vaccine regiments and experimental set-ups. The single cell RNA-seq technique used here is not only able to identify the memory B cell population of interest, but also five additional B cell subsets based on subset-specific markers implying both T cell dependent and T cell independent formation. More importantly, it samples deeply enough to pick up several highly similar memory B cells based on the usage of the same VH and VK combinations that appear reproducibly even between mice. In combination with the high expression levels of the Mki67 proliferation marker, this suggests a current clonal expansion of the switched MBC population against fentanyl. When the BCRs of the switched MBC population are cloned as mAb, every single one of these exhibits very high affinity to fentanyl in the picomolar range. Additionally, we were able to detect two memory B cells with nearly identical BCR sequences except for a couple of point mutations (FenAb196 and FenAb208), which also share some somatic hypermutations compared to the germline sequence, strongly suggesting that these cells originate from the same progenitor cell; another line of evidence suggesting that the VAST platform is indeed able to induce true immunological memory.

Although anti-fentanyl mAb have been described to effectively protect from fentanyl challenge before (Smith *et al*., 2019; Baehr *et al*., 2020; Ban *et al*., 2021), the modality of fentanyl binding of these antibodies has yet to be elucidated. Interestingly, all of our crystallized antibodies bind fentanyl in a strikingly deep and narrow pocket. This pocket is most likely responsible for the high specificity of the antibodies as only very similar small and elongated structures would be able to fit. This finding is particularly important considering the implications an anti-fentanyl vaccine could have for the clinic. If the antibodies cross-react with other common clinically relevant opioid painkillers or anesthetics, the approach could prove problematic. On the contrary, antibodies elicited by this vaccination scheme show, so far, no evidence of being “pan-opioid” and thereby remain promising preclinical candidates. For instance, antibodies previously raised against the fentanyl-hapten used in our studies showed selectivity for fentanyl, sufentanil, and acetylfentanyl but not methadone, buprenorphine, naloxone, naltrexone and critical care medications such as anesthetics (Robinson *et al*., 2020).

In conclusion, the VAST platform can be used both as a vaccine itself and as a platform to identify highly-protective monoclonal antibodies for direct therapeutic or research applications. This epitope focused platform may serve as a tool to help unlock certain fields of therapeutic antibody development that have yet to be successfully explored. Future studies will address this hypothesis that the VAST platform can elicit high-affinity antibodies for a wide variety of different antigen classes, many of which are already underway (Figure 1C).

## Acknowledgements

We acknowledge synchrotron time at the Paul Scherrer Institut, Villingen, Switzerland (SLS, beamline PXIII, Vincent Olieric and colleagues). We acknowledge the DKFZ Flow Cytometry core facility and the DKFZ Single-Cell Open Lab (scOpenLab) for performing key workflows towards the identification of antibody sequences. We thank Dustin Hicks and Jenny Vigliaturo (University of Minnesota) for key experimental assistance. We thank Vanessa Hofmann for mass spectrometric analysis. This project was funded by grants from the NIH (R01AI097127 to F.N.P and 1R43DA052960-01 to J.P.V) and funds from the Helmholtz Foundation (to F.N.P and C.E.S). We also acknowledge early contributions by P. Stavropoulos and J. Pinger, who helped initiate the work presented here.

## Author Contributions

C.E.S. and F.N.P. conceived and initiated the project. Immunological Work

A.B. and A.K.M. generated fen-G4 and fen-sort.

H.H. produced the sortaggable trypanosome lines.

A.S., G.T., H.H., and J.P.V. conducted sortagging experiments.

G.T. and H.H. performed ELISA assays.

F.N.P., G.T., H.H., and S.K. conducted immunization experiments and isolated immunological tissues.

F.N.P., G.T., and S.R. and conducted fentanyl challenges and behavioral assays.

M.P. coordinated and analyzed mass spectrometry. Bioinformatic Derivation of Antibodies

P.V. conceived and constructed the bioinformatic pipeline and analyzed the SmartSeq data with assistance from G.T. and F.N.P., deriving antibody sequences.

Antibody Characterization

M.v.S. cloned and produced Fabs and Y.K. cloned and produced Mabs.

C.B. performed ELISA and Octet assays.

A.S. performed and analyzed ITC.

F.N.P. and G.T. performed and analyzed passive therapy experiments

M.B. performed mass spectrometry Crystallography

M.v.S. crystallized Fab-fentanyl complexes.

J.P.Z. and M.v.S. prepared crystals for beamline data collection, collected the data, and solved the structures.

C.E.S., J.P.Z., and M.v.S analyzed the structural data.

C.E.S., C.L.M., F.N.P, G.T., J.P.V and M.P. coordinated the project. C.E.S., G.T. and J.P.V. wrote the manuscript.

## Competing interests

G.T. reports being a shareholder of Panosome GmbH and Hepione Therapeutics and having patents planned, pending, or issued broadly relevant to the work. J.P.V reports being the C.S.O. and shareholder of Panosome GmbH and Hepione Therapeutics and having patents planned, pending, or issued broadly relevant to the work. C.E.S reports being shareholder and managing director of Panosome GmbH and CEO of Hepione Therapeutics and having patents pending or issued broadly relevant to the work. F.N.P reports being shareholder and managing director of Panosome GmbH and shareholder of Hepione Therapeutics and having patents pending or issued broadly relevant to the work. P.V. reports being an employee of Panosome GmbH and having patents planned, pending, or issued broadly relevant to the work. S.R. reports being an employee of Panosome GmbH. S.K. reports being a shareholder of Panosome GmbH and Hepione Therapeutics. M.v.S., J.P.Z., Y.K., A.S., A.K.M, A.B. and H.H. report having patents planned, pending, or issued broadly relevant to the work. M.P. and C.B. have patents pending or planned related to fentanyl haptens, conjugates, and monoclonal antibodies against fentanyl-like compounds.

## Star Methods

### Lead Contact

Further information and requests for resources and reagents should be directed to and will be fulfilled by the lead contact, Dr. Joseph P. Verdi.

### Data and materials availability

#### Materials Availability

All newly generated materials produced by these studies are available for distribution under certain conditions. The VAST platform, and all key reagents associated with it or derived from its use in animals, are licensed for commercial purposes to Panosome GmbH and/or Hepione Therapeutics. These companies must authorize any material distributions to third parties and be privy to the associated material transfer agreements established therein.

#### Data and Code Availability

The following structure files are available from the Protein Databank: 7QT0, 7QT2, 7QT3, and 7QT4. FASTq files will be made available upon request.

### Experimental Models and Subject Details Mice

Female C57BL/6J mice (6-8-weeks old) were purchased from Janvier and handled in accordance with the German Animal Protection Law (§8 Tierschutzgesetz) and approved by the Regierungspräsidium Karlsruhe, Germany (project numbers Aktenzeichen 35-9185.81/G-285/18). Mice were randomly assigned to experimental groups which were kept in separate cages.

### T. brucei brucei parasites

All trypanosome cell lines used in this study were bloodstream-form trypanosomes ultimately derived from the Lister 427 cell line (Wirtz *et al*., 1999). Trypanosomes were cultivated in HMI-9 medium with 10% fetal bovine serum at 37 °C and 5% CO_2_. All trypanosomes used in this study expressed a modified version of VSG3 (S317A). The S317A variant was genetically engineered to lack an o-linked glycan present at the 317th residue of the wild-type counterpart. The newly engineered variant (Srt-VSG3) additionally contains the sortagging motif AA on the N-terminus of the VSG that has been extended above the top of the surface coat through the addition of a linker region consisting of 3 Gly-Gly-Gly-Gly-Ser motifs. They were either expressing (GPI-PLC WT) or lacking GPI-PLC (GPI-PLC -/-). The GPI-PLC WT cells used were the 2T1 cell line (Alsford and Horn, 2008), while the GPI-PLC -/- counterparts were derived from the 2T1s in-house using knockout vectors kindly provided by Dr. M. Carrington. GPI-PLC -/- cells lack the ability to shed their VSG and thus stay intact during UV irradiation.

### Human embryonic kidney cells used for antibody production

Human Embryonic Kidney (HEK) “FreeStyle” 293F cells (ThermoFisher Scientific R79007) were grown in tissue culture flasks in FreeStyle^TM^ 293 Expression Medium (ThermoFisher Scientific 12338018). Cells were incubated at 37 °C with 5-8% CO_2_ on a shaking platform operating at 120-130 rpm.

## Method details

### Experimental design

Female mice were used in all experiments and were randomly assigned to the various experimental groups. All experiments were performed using age-matched mice from 6 to 8 weeks old at the start of the experiment. Similar vaccination experiments to those shown in Figure 2 were performed extensively with similar results. The single-cell sequencing approach to B cell repertoire analysis was performed after consulting the literature for known markers and ensuring that population identification was robust. Statistical analyses are detailed throughout the manuscript.

### Synthesis of fentanyl derivatives

Lithium 5-oxo-5-((2-(4-(*N*-phenylpropionamido)piperidin-1-yl)ethyl)amino)pentanoate (**3**): To a solution of the bis trifluoroacetate salt of amine **1** (compound **5** in Raleigh et al. (Raleigh *et al*., 2019)) (16.513 g, 32.8 mol) in anhydrous CH_2_Cl_2_ (250 ml) was added pyridine (15.9 ml, 197 mmol, 6.0 equiv.) followed by glutaric acid monomethyl ester chloride (4.54 ml, 32.8 mmol, 1.0 equiv.) at 0 °C under argon. After addition, the cooling bath was removed and after 16 h, the reaction was diluted with CH_2_Cl_2_ (150 ml), washed with saturated aqueous NaHCO_3_ (3 x 200 ml) and brine (100 ml). The organic layer was dried (MgSO_4_), filtered, and concentrated in vacuo. The product was purified by column chromatography (5% to 7.5% gradient of MeOH in CH_2_Cl_2_) to give ester **2** (9.79 g, 74%) as a colorless oil. ^1^H NMR (400 MHz, CDCl_3_, CHCl_3_ referenced to 7.26 ppm) δ 7.40–7.32 (m, 3H), 7.06–7.02 (m, 2H), 6.00 (br s, 1H), 4.58 (tt, *J* = 12.2, 3.9 Hz, 1H), 3.58 (s, 3H), 3.22 (q, *J* = 5.8 Hz, 2H), 2.83 (m, 2H), 2.36 (t, *J* = 6.0 Hz, 2H), 2.28 (t, *J* = 7.2 Hz, 2H), 2.12 (t, 7.2 Hz, 2H), 2.07 (m, 2H), 1.89–1.81 (m, 4H), 1.76–1.70 (m, 2H), 1.32 (m, 2H), 0.95 (t, *J* = 7.5 Hz, 3H) ppm; ^13^C NMR (101 MHz, CDCl3, CHCl3 referenced to 77.16 ppm) δ 173.7, 173.6, 172.1, 139.0, 130.4, 129.4, 128.4, 56.7, 53.0, 52.3, 51.6, 36.2, 35.4, 33.1, 30.5, 28.6, 20.9, 9.7 ppm; LC/MS (ESI^+^) *m/z* = 404.2. To a solution of methyl ester **2** (9.65 g, 23.9 mmol) in MeOH (120 ml) was added a solution of LiOH (1.72 g, 71.7 mmol, 3.0 equiv.) in H_2_O (30 ml) at room temperature. After 22 h, the reaction was partially concentrated in vacuo and completely dried via lyophilization to give unpurified lithium carboxylate **3** as a white powder (9.06 g, ∼85%). ^1^H NMR (400 MHz, D_2_O) δ 7.53–7.47 (m, 3H), 7.25–7.21 (m, 2H), 4.45 (tt, *J* = 12.2, 3.9 Hz, 1H), 3.25 (t, *J* = 7Hz, 2H), 2.91 (m, 2H), 2.45 (t, *J* = 7.0 Hz, 2H), 2.25–2.11 (m, 6H), 1.97 (q, *J* = 7.6 Hz, 2H), 1.82–1.73 (m, 4H), 1.32 (dq, *J* = 12.5, 3.5, 2H), 0.92 (t, *J* = 7.5 Hz, 3H) ppm; ^13^C NMR (101 MHz, D_2_O, referenced to external TSP = 0.0 ppm) δ 185.3, 179.8, 179.1, 140.7, 132.6, 132.4, 131.8, 58.4, 55.7, 54.9, 39.6, 39.1, 38.3, 32.0, 31.2, 25.2, 12.1 ppm; LC/MS (ESI^-^) *m/z* = 388.2. This material was used without further purification for solid phase synthesis.

### Solid-phase peptide synthesis procedure definitions

*WASH:* The resin is suspended in DMF (3 ml) with agitation for 2 min, followed by disposal of the solvent. This process is repeated two more times for a total of three washes.

*Fmoc-OFF:* The resin is suspended in a 20% solution of piperidine in DMF (3 ml) with agitation for 5 min, followed by disposal of the solution. This process is repeated one more time, but with an incubation time of 7 min.

*COUPLING* (name of coupling partner): A DMF solution (3 ml) containing “name of coupling partner” (4.0 molar equivalents relative to the initial resin loading), PyBOP (4.0 equiv.), and *i*-Pr_2_NEt (8.0 equiv) is prepared and pre-activated for 10 min. The resin is then suspended and agitated in this cocktail for 1 h (unless otherwise noted), followed by disposal of the solution.

Fen-G4: Briefly, fen-G4 was prepared by solid-phase peptide synthesis starting with a glycine-labeled resin, followed by coupling to Fmoc-GGG-OH (BACHEM) and then compound **3**. In detail, a syringe (5 ml) for peptide synthesis (frit made of PE) was charged with pre-loaded Fmoc-G-SASRIN resin (417 mg, loading: 0.72 mmol/g, BACHEM) and swelled for 1 h in DMF (3 ml). The synthesis procedure then proceeded according to the above definitions: *WASH, Fmoc-OFF, COUPLING* (Fmoc-GGG-OH, BACHEM, for 2 h), *WASH, Fmoc-OFF, WASH, COUPLING* (compound **3**), *WASH*. The resin was then washed (2 min) with CH_2_Cl_2_ (3 ml) three times. For cleavage, the resin was treated (5 min) with 80% trifluoroacetic (TFA) in water (3 ml) twice, wherein the resin beads became dark purple. The two solutions were combined and concentrated in vacuo. This residue was dissolved in water (ca. 25 ml) and lyophilized. The product was purified via reverse-phase chromatography (C18 silica, 1 to 40% gradient of MeCN in H_2_O) and then lyophilized with a few drops of added TFA to obtain fen-G4 as a trifluoroacetate salt (49 mg, 0.067 mmol, 22%). ^1^H NMR (400 MHz, D_2_O, HOD referenced to 3.79 ppm) δ 7.57–7.48 (m, 3H), 7.29–7.23 (m, 2H), ∼4.75 (1H, obscured by solvent, confirmed by HSQC), 3.99 (s, 6H), 3.93 (s, 2H), 3.66 (d, *J* = 12.2 Hz, 2H), 3.54 (m, 2H), 3.23 (m, 2H), 3.16 (t, *J* = 12.4 Hz, 2H), 2.33–2.22 (m, 4H), 2.13 (d, *J* = 13.8 Hz, 2H), 2.01 (q, *J* = 7.4 Hz, 2H), 1.84 (app pent, *J* = 7.2 Hz, 2H), 1.64 (q, *J* = 12.5 Hz, 2H), 0.94 (t, 7.2 Hz, 3H) ppm; ^13^C NMR (101 MHz, D_2_O, referenced to internal MeOD = 49.5 ppm) d 178.9, 178.2, 177.9, 174.7, 173.8, 173.5, 173.2, 164.4 (q, ^2^*J*_CF_ = 36.4 Hz), 138.5, 131.2 (2C), 130.7, 117.7 (q, ^1^*J*_CF_ = 291.3 Hz), 57.4, 53.8, 51.2, 43.9 (2C), 43.7, 42.5, 35.8 (2C), 35.6, 29.7, 28.8, 22.4, 10.5 ppm; HRMS (ESI^-^) *m/z* calcd for [C_29_H_42_N_7_O_8_^-^]: 616.3100; found: 616.3104.

Sortaggable Fentanyl (fen-sort): Briefly, fen-sort was prepared by solid-phase peptide synthesis starting with a glycine-labeled resin, followed by sequential coupling to Fmoc-protected G, T, S, P, L, S, G, G, G (all from BACHEM), and then compound **3**. In detail, a syringe (5 ml) for peptide synthesis (frit made of PE) was charged with pre-loaded Fmoc-G-SASRIN resin (417 mg, loading: 0.72 mmol/g, BACHEM) and swelled for 1 h in DMF (3 ml). The synthesis procedure then proceeded according to the above definitions: *WASH, Fmoc-OFF, COUPLING* (Fmoc-G-OH), *WASH, Fmoc-OFF, WASH, COUPLING* (Fmoc-T(*t*-Bu)-OH), *WASH, Fmoc-OFF, WASH, COUPLING* (Fmoc-S(*t*-Bu)-OH; Note: collidine was used instead of *i*-Pr_2_NEt in this step), *WASH, Fmoc-OFF, WASH, COUPLING* (Fmoc-P-OH), *WASH, Fmoc-OFF, WASH, COUPLING* (Fmoc-L-OH), *WASH, Fmoc-OFF, WASH, COUPLING* (Fmoc-S(*t*-Bu)-OH; Note: collidine was used instead of *i*-Pr_2_NEt in this step), *WASH, Fmoc-OFF, WASH, COUPLING* (Fmoc-G-OH), *WASH, Fmoc-OFF, WASH, COUPLING* (Fmoc-G-OH), *WASH, Fmoc-OFF, WASH, COUPLING* (Fmoc-G-OH), *WASH, Fmoc-OFF, WASH, COUPLING* (compound **3**), repeat *COUPLING* (compound **3**), *WASH*. The resin was then washed (2 min) with CH_2_Cl_2_ (3 ml) three times. For cleavage the resin was treated (5 min) with a cocktail of 90% TFA, 5% water and 5% triisopropylsilane (3 ml) three times, during which the resin beads became dark purple. The three washings were combined and stirred for 1 h, and then added dropwise to cold Et_2_O (50 ml). The obtained precipitate was filtered and washed thoroughly with cold Et_2_O. The solid residue was taken up in water and lyophilized. The product was purified via reverse-phase MPLC (C18 silica, 1 to 40% gradient of MeCN in H_2_O) and then lyophilized with a few drops of added TFA to obtain fen-sort as a trifluoroacetate salt (99 mg, 0.078 mmol, 26%). ^1^H NMR (400 MHz, D_2_O, HOD referenced to 3.79 ppm) d 7.56–7.49 (m, 3H), 7.29–7.24 (m, 2H), ∼4.75 (1H, obscured by solvent, confirmed by HSQC), 4.65 (dd, *J* = 9.9, 4.5 Hz, 1H), 4.52–4.44 (m, 3H), 4.41 (d, *J* = 3.8 Hz, 1H), 4.31 (dq, *J* = 6.4, 3.8 Hz, 1H), 4.03–3.97 (m, 7H), 3.96–3.91 (m, 3H), 3.90–3.81 (m, 4H), 3.70–3.63 (m, 3H), 3.55 (d, *J* = 6.2 Hz, 2H), 3.24 (d, *J* = 6.2 Hz, 2H), 3.20–3.12 (m, 2H), 2.35–2.23 (m, 5H), 2.14 (d, *J* = 13.8 Hz, 2H), 2.08–1.90 (m, 5H), 1.85 (sept, *J* = 7.3 Hz, 2H), 1.72–1.55 (m, 5H), 1.22 (d, *J* = 6.4 Hz, 3H), 0.97–0.90 (m, 9H) ppm; ^13^C NMR (101 MHz, D_2_O, referenced to internal MeOD = 49.5 ppm) d 178.9, 178.2, 177.9, 175.8, 174.5, 174.1, 173.8 (3C), 173.5, 173.1, 172.8 (2C), 164.3 (q, ^2^*J*_CF_ = 35.6 Hz), 138.5, 131.2 (2C), 130.7, 117.8 (q, ^1^*J*_CF_ = 291.6 Hz), 68.4, 62.6, 62.3, 61.9, 60.5, 57.4, 57.0, 56.8, 53.8, 51.8, 51.2, 49.3, 43.9 (2C), 43.8 (2C), 42.4, 40.5, 35.8 (2C), 35.6, 30.8, 29.7, 28.8, 26.1, 25.8, 23.9, 22.4, 22.1, 20.2, 10.5 ppm; HRMS (ESI^-^) *m/z* calcd. for [C_52_H_80_N_13_O_17_^-^]: 1158.5801; found: 1158.5810.

### Purification of Srt-VSG3 protein

Srt-VSG3 protein was purified from *T. brucei* PLC WT expressing Srt-VSG3 as described elsewhere (Pinger, Chowdhury and Papavasiliou, 2017). Briefly, cells were cultured *in vitro* in HMI-9 media to a density of 4×10^6^ cells/ml. Cells were pelleted, lysed in 0.2 mM ZnCl_2_ + HALT protease inhibitor and then the lysis mixture was centrifuged at 10,000xg for 10 minutes. The pellet which contained the membrane material was resuspended in pre-warmed (40 °C) 20 mM HEPES pH 7.5 with 150 mM sodium chloride (NaCl), enabling the activation of endogenous lipases and resulting in the efficient release of surface VSG protein from the membrane. The membranous material was then pelleted two more times, while supernatants (containing soluble VSG) were collected. Supernatants were loaded onto an anion-exchange column (Q-Sepharose Fast-Flow, GE Healthcare), which had been equilibrated with 20 mM HEPES buffer with 150 mM NaCl (the VSG does not bind to the column, while contaminating proteins are trapped). Srt-VSG3 was then separated from remaining contaminants and aggregation products via a gel filtration column (Superdex 200, GE Healthcare) equilibrated in 20 mM HEPES buffer with 150 mM NaCl. Aliquots from the gel filtration runs were analyzed on SDS–PAGE for visual inspection (Suppl. Figure 1E).

### Sortagging of soluble Srt-VSG3

Sortase A was expressed and purified as described (Pinger, Chowdhury and Papavasiliou, 2017). Sortagging solutions containing 100 µM purified Sortase A and 300 µM of fentanyl-LPSTGG in PBS or 20 mM HEPES, 150 mM NaCl, pH 8.0 were incubated on ice for 30 min. Purified Srt-VSG3 protein was added to the sortagging solution at a concentration of not higher than 2 mg/ml and incubated for 2 h at 37 °C while gently shaking and then at 4 °C rotating overnight. Fentanyl-sortagged Srt-VSG3 protein was re-purified via gel filtration (Superdex 200, GE Healthcare), concentrated to not higher than 2 mg/ml, and flash frozen for storage at −80 °C until further use.

### Sortagging of intact *T. brucei brucei*

In order to avoid the naturally occurring shedding of the VSG coat, a protection mechanism of trypanosomes from antibody attachment in the host, Srt-VSG3 PLC-/- *T. brucei* was used throughout the whole paper. Srt-VSG3 PLC-/- *T. brucei* expresses VSG3 on the surface and lacks the enzyme PLC which is responsible for the natural VSG shedding. For sortagging whole trypanosomes, the trypanosomes were incubated with a sortagging solution containing 100 µM purified Sortase A and 300 µM of fen-sort in HMI-9 media on ice for 30 min. PLC -/- trypanosomes expressing Srt-VSG3 were pelleted, resuspended in the sortagging solution at a concentration of 10^8^ cells/ml, and incubated for 2 h at 4 °C on an inversion rotator. Cells were then pelleted, washed once with HMI-9 media, and pelleted again before resuspension in HMI-9 or staining solution containing an anti-fentanyl monoclonal antibody (provided by M. Pravetoni, University of Minnesota) conjugated to FITC (Abcam, ab102884) for analyzing sortagging success. Trypanosomes were immediately analyzed using a BD FACS Calibur and FlowJo software.

### UV irradiation of sortagged *T. brucei brucei*

After sortagging with fentanyl, the trypanosomes were pelleted, washed with irradiation buffer (PBS-glucose (10 g/l)) and resuspended in irradiation buffer to a density of 10^8^ cells/ml. A volume of 1 ml of this resuspension was aliquoted into each well of a non-treated six-well tissue culture plate. Plates were UV-irradiated for four times in 1 min intervals using a FB-UVXL-1000 UV crosslinker. Plates were swirled between 1 min intervals to ensure equal irradiation of trypanosomes. Complete irradiation was confirmed using a microscope and cells were counted and diluted for immunization.

### Mouse immunizations

Five 6-8 weeks old female C57BL/6J mice per group were primed at day 0 and 14 or 30 with 5×10^6^ intact UV-irradiated trypanosomes, either sortagged or not sortagged with fentanyl, and without adjuvant via subcutaneous injection. The mice then received a booster of 100 µg of soluble Srt-VSG3, either sortagged or not sortagged with fentanyl (without adjuvant) at days 42 or 60 and 70 or 90. Serum samples were taken 2 days before and one week after injection.

### Serum ELISA

The fen-G4 peptide was conjugated to BSA via EDC crosslinking as a heterologous carrier protein and used to coat 96-well ELISA plates at 10 µg/ml overnight at 4 °C. Similarly, plates were coated with FPLC-purified Srt-VSG3 protein at 5 µg/well. Plates were blocked for 1.5 h at RT with 4% BSA in PBS. Coated plates were incubated with the serum for 1.5 h at RT. As a control, an anti-VSG3 monoclonal antibody or an anti-fentanyl monoclonal antibody were used at a starting concentration of 1 µg/ml or 5 µg/ml, respectively, and diluted 4-fold. Bound antibodies were detected by goat anti-mouse IgG secondary antibody coupled to horse-radish-peroxidase (HRP) (Jackson Immuno Research) diluted 1:3,000 in 1% BSA in PBS which was then detected using an ABTS substrate solution complimented with H_2_O_2_ (Roche). The optical density (OD) at 405 nm was determined on an M1000Pro plate reader (Tecan) after 40 min. Graphs were created, area under curve (AUC) values were calculated and then normalized to the respective control antibody using GraphPad Prism 6.07.

### Antinociception

Analgesic activity was tested by using the hotplate antinociception assay as described by Cox and Weinstock (Cox and Weinstock, 1964). Baseline antinociception of the mice was measured by placing them on a hot plate at 54 °C and measuring the latency to a response (in seconds) like flipping or licking a paw or jumping. In order to avoid tissue damage the test was aborted after 60 s without a reaction. Then mice were injected subcutaneously with an initial dose of 50 µg/kg fentanyl. After 30 min antinociception was measured. Mice received another dose of 50 µg/kg (cumulative dose of 100 µg/kg) and 15 min later antinociception was measured again. 15 min after that mice received a single dose of naloxone (100 µg/kg), which reverses the effects of fentanyl, antinociception was measured again. The effect of fentanyl and naloxone was measured as latency to response as well as the percentage maximum possible effect (% MPE). MPE was calculated as the post-test latency minus the pretest latency divided by the maximum time (60 seconds) minus the pretest latency times 100 (Raleigh *et al*., 2019).

### Straub-Tail Reaction

The Straub tail reaction is a dorsiflexion of the tail that is often almost vertical to the orientation of the body or curling back over the animal and stereotyped walking behavior (Bilbey, Salem and Grossmann, 1960). This phenomenon was first described as a response to opiates in mice (Straub (1911) cited by (Bilbey, Salem and Grossmann, 1960)), and is thought to be mediated by activation of the opioid receptor system because opioid receptor antagonists such as naloxone block the phenomenon (Aceto, McKean and Pearl, 1969; Nath *et al*., 1994; Zarrindast, Alaei-Nia and Shafizadeh, 2001). The Straub-Tail reaction was recorded as present (+) or absent (-).

### Analysis of fentanyl concentrations in tissue: extraction and LC-MS/MS conditions

#### Preparation of brain samples

Brain tissue was homogenized using Agilent ceramic beads for 4 min with a Beadblaster 24 homogenizer (Benchmark Scientific, Sayreville, NJ) at 7 m/s, then centrifuged briefly to reduce bubbles. The homogenate was transferred to a cryogenic tube and placed at −20°C until time of extraction.

#### Sample extraction

Extraction of serum, brain homogenate, and standards was performed at 4 °C. For standards, 20 µL of stock calibrator solution was added to 180 µL of fetal bovine serum. 200 µL of sample was used for extraction with 20 µL of internal standard solution added to all samples. 600 µL of cold LC-MS grade Acetonitrile was added to 1.6 ml microcentrifuge tubes to precipitate proteins and then centrifuged at 8,609xg for 10 min. Supernatant was transferred to a 2 ml 96-well collection plate, evaporated to 200 µL on a Mini Vap (Porvair Sciences, Wrexham, UK) and then diluted 1:4 with 2% phosphoric acid. Extraction was performed using Bond Elut 96, Plexa PCX, 1 ml, 30 mg (Agilent, Santa Clara, CA). Cartridges were first washed with 500 µL methanol followed by 500 µl water and then samples were loaded onto the plate. The plate was washed in series, first using 600 µl of 2% formic acid followed by 600 µL 1:1 methanol/acetonitrile. The plate is dried on a Positive Pressure Manifold (Agilent, Santa Clara, CA), placed above fresh round bottom 1 ml 96-well collection plate to elute samples using 750 µL of 5% ammonium hydroxide in 1:1 methanol/acetonitrile, and dried on the Mini Vap. Samples were reconstituted in 200 µL LC-MS grade water, 0.1% ammonium formate, 0.01% LC-MS grade formic acid (mobile phase A).

#### LC-MS/MS conditions

Two microliters of sample were injected onto a reversed phase Agilent (Santa Clara, CA) Poroshell 120 SB-C18 column (2.1 mm x 50 mm 2.7 μm) column. The LC-MS/MS system consisted of an Agilent G6470A triple quadrupole with an Infinity II 1290 G7116B Multicolumn Thermostat, G7120A High Speed Quad Pumps, G7267B Multisampler. The samples were kept at 4 °C during injection. Gradient elution was performed with a mixture of mobile phase A and methanol, 0.01% formic acid (mobile phase B) as follows: 0-0.5 min 5% mobile phase B, 0.5-2.25 min 15 → 50% mobile phase B, 2.25-4.0 min 50 → 95% mobile phase B, 4.0-6.0 95% mobile phase B. The flow rate was kept at a constant 0.40 ml/min and the total run time was 6 min. Electrospray ionization was achieved by Agilent Jet Stream high sensitivity ion source in the positive ion mode. Instrument settings were: gas temperature 325 °C, gas flow 9 L/min, nebulizer pressure 40 psi, sheath gas temperature 380 °C, sheath gas flow 10 L/min, capillary 2500 V, and pos nozzle 0V. Data acquisition and peak integration were interfaced to a computer workstation using Mass Hunter (Tokyo, Japan). High-performance liquid chromatography-tandem mass spectrometry (LC-MS/MS) and their specific mode was used for the mass spectrometric analysis to identify the appropriate ions to monitor *m/z*: fentanyl 337.2 – 188.1, secondary 337.2 – 105.1, fentanyl-d5 342.3 – 188.1, secondary 342.3 – 105.1. The analysis of extracted serum samples from the mice after dosing (26.25 and 3.75 mg/kg) as well as the controls were also measured by applying LC-MS/MS and fentanyl concentrations were determined after appropriate sample preparation following the above-described method.

#### Flow cytometry and single cell sorting

Fentanyl-reactive B cells were analyzed on a LSR II instrument and isolated using an ARIA II cell sorter (BD Bioscience). Cells were stained using fentanyl-PE (1:1000; 1 µM stock) and a decoy-PE-AF647 (1:50) (Pravetoni *et al*., 2014; Laudenbach *et al*., 2015; Baruffaldi *et al*., 2018) and the following antibodies: rat anti-mouse CD19-BV412 (1:100, Biolegend), rat anti-mouse IgG1-BV650 (1:100, Biolegend), rat anti-mouse CD138-BV510 (1:300, Biolegend), rat anti-mouse GL-7-FITC (1:1000, BD Pharmingen), rat anti-mouse CD38-PE-Cy7 (1:400, Biolegend), goat anti-mouse IgM-biotin (1:400, Jackson), rat anti-mouse IgD-APC-Cy7 (1:1000, Biolegend), streptavidin-BV785 (1:400, Biolegend). LIVE/DEAD™ Fixable Blue Dead Cell Stain Kit (1:1000, Invitrogen) was included in all stainings to exclude dead cells. The data were analyzed using FlowJo v10 software. For single-cell sorting, fentanyl-reactive B cells were defined as live fentanyl+decoy-CD19+ and sorted into 384-well plates containing lysis buffer using an ARIA II cell sorter (BD Bioscience and using the index-sort function).

### SMART-seq 2.5 library preparations

Single-cell RNA sequencing was performed following the SMART-seq 2.5 library preparation protocol as described in Picelli et al. (Picelli *et al*., 2014), and modified by the DKFZ scOpenLab in Bioquant, Heidelberg. Plate preparation was assisted by the BRAVO Agilent and Mosquito LV liquid handling platforms. Quality control was performed on the TapeStation provided by Agilent Technology, using a D5000 Tape. Single cell library preparation was performed using the Nextera XT DNA SMP Prep Kit (96 SMP). DNA was cleaned-up using a magnetic size selection. The samples were finally quantified and assessed on a D1000 TapeStation.

### Single cell RNA Sequencing

Single cell RNA sequencing was performed on the Illumina sequencing platform of the DKFZ Genomics and Proteomic sequencing facility. The samples were pooled and delivered as multiplexed. Next-generation sequencing (NGS) of 75 bp single reads was performed using the NextSeq 550 Sequencing System. Quality control was performed before and after the sequencing process. The quality of the raw reads was assessed using fastqc and MultiQC (Ewels *et al*., 2016). Adapter (Nextera transposase sequence) and quality trimming (Phred score cutoff of 20, overlap of 3 bp) was performed with the TrimGalore version: 0.6.4_dev (Cutadapt version: 1.18) (Krueger, 2015), keeping the sequencing reads with a length of at least 36 bps. Genome reference-based alignment was performed using the slice-aware STAR alignment tool with the default parameters with Release M25 (GENCODE-GRCm38.p6) of the mouse genome. The reference genome was indexed with a bp overhang of 100-1. The generated bam files were sorted using Samtools based on the chromosome coordinates. Unmapped reads, PCR duplicates, and reads with an alignment p.score <20 were removed by filtering the sorted bam files.

### Transcriptome analysis

Downstream preprocessing analysis was performed separately for the sorted plates in the R programming language using the Bioconductor packages SingleCellExperiment (1.16.0), scater (1.22), and scran (1.22.1). The count matrices with only the exonic sequences were generated using the *summarizeOverlaps* function of the GenomicAlignments package (1.30 version -(Lawrence *et al*., 2013)). Gene annotation was performed with the *makeTxDbFromGFF* function of the GenomicFeatures package (1.46.1), the AnnotationDbi 1.52.0 and the org.Mm.eg.db 3.12.0 package. The mitochondrial genes were identified by extracting the genomic coordinates according to the *TxDb.Mmusculus.UCSC.mm10.ensGene* data set.

### Quality control, normalization, and feature selection

Outlier cells with high mitochondrial percentage, low library numbers and low unique features were identified and removed based on a median absolute deviation (MAD) = 3. Additionally, gene filtering was applied by removing genes with <3 reads in <5 cells. Further, we also eliminated transcripts that map to the VDJ immunoglobulin genes, by keeping the reads aligning to the constant regions of the heavy and light chains. The gene expression was normalized by deconvolution by applying the *quickCluster* and *ComputeSumFactors* functions of scran. Next, we computed the log-transformed normalized expression values. Feature selection was performed with *modelGeneVar*, which models the variance of the log-expression profiles for each gene on a fitted mean-variance trend. Genes with a variance of >0 and FDR <0.1 are maintained (∼13,000 for each plate).

### Data integration

The sorted plates (262 and 288 cells, respectively, for each plate) were integrated based on non-negative matrix factorization using the rLiger package (1.0.0), by asserting all the genes that are either shared or unique for each plate and have a variance >0.1 and an alpha threshold of 0.05 (Welch *et al*., 2019). The lambda and kappa values were fixed to 5 and 20 respectively. Genes were normalized and scaled, and the cells were clustered using a resolution of 0.9. Finally, Uniform Manifold Approximation and Projection (UMAP) was applied by computing the ‘cosine’ distance of 30 neighboring cells with a minimum distance of 0.1.

### Differential expression and marker gene identification

A first characterization of the B cell subclusters was performed by using a set of known B cell markers. Umap visualizations of the joint expression of 3 known memory B cell markers were created using the Nebulosa package (1.4 version - (Alquicira-Hernandez and Powell, 2021)). Differential gene expression between clusters was performed with the MAST algorithm (Finak *et al*., 2015) integrated in the Seurat:4 package (Hao *et al*., 2021) after transforming the liger object into a Seurat object of 550 cells, using the default parameters. For the subcluster characterized as “switched memory B cells”, more stringent parameters were applied (at least <0.5 LFC and the presence of the markers in at least 30% of the cells) to reassure the identification of robust marker genes. For visualization purposes, the z-score of the top 10 genes from each subcluster with a positive log fold change along with several known B cell markers were plotted together. Using the *pheatmap* package hierarchical clustering between both the selected genes (rows) and the identified subclusters (columns) was applied. Columns were clustered based on the “correlation” distance and the “ward.D” method, while rows were based on the “euclidean” distance.

### BCR sequencing analysis

The trimmed reads of the raw fastq files were analyzed using the BASIC vdj assemble package (version 1.5.0) (Canzar *et al*., 2017). Annotation of the VDJ segments was performed with the Igblast (v1.16.0) using a database of V(D)J genes from IMGT as a reference while blast (version 2.11.0) was used for the identification of the constant chains. Downstream analysis was performed in R, using the Change-O - Repertoire clonal assignment toolkit from the “Immcantation” portal (version 1.0.0 (Gupta *et al*., 2015)) and Shazam (version 1.1.0) for the mutational load analysis (Gupta *et al*., 2015).

### Plots and figures

The plots shown in Figure 4 as a part of the sequencing analyses were created using ggplot2 (version 3.3.5), gridExtra (v 2.3) and ggrepel (v 0.9.1), joint density plots with Nebulosa (v 1.4), heatmap with pheatmap (1.0.1.) and circos plots were formed using the circlize package (version 0.4.13). The images represented throughout the rest of the manuscript were generated using Adobe Photoshop and Adobe Illustrator after rendering plots and tables in GraphPad Prism, Microsoft Powerpoint, Microsoft Excel, and/or FlowJo. Structures and molecular surfaces are illustrated with CCP4mg52 (McNicholas et al., 2011; de Beer et al., 2014, Schrödinger, 2015). The images shown in Figures 1B and Suppl. Figure 1F were generated with the assistance of BioRender.

### Monoclonal antibody cloning

For IgGs, VDJ regions were synthesized via Twist Biosciences with flanking restriction sites matching the expression vectors taken from Tiller et al. (Tiller *et al*., 2008). Heavy chain VDJ regions were flanked with a 5’ AgeI restriction site and 3’ SalI restriction site. Light chain VDJ regions were flanked with a 5’ AgeI restriction site and 3’ BsiWI restriction site. Heavy and light chain vectors and VDJ region-containing plasmids were digested as per Tiller et al. using enzymes obtained from NEB. Linear inserts were ligated into the appropriate linear vector using T4 DNA Ligase (ThermoFisher Scientific EL0014). The resulting expression vectors encode antibodies under the control of a CMV promoter. For Fabs, VDJ regions of the Fab heavy chain FenAB136, 609, 709 and FenAB136 light chain were synthesized commercially (Thermo Fisher Scientific, GeneArt), while the VDJ regions of FenAB208 were cloned out of the IgG vector templates. Mammalian expression vectors containing the various Fab heavy chain genes, as well as those containing light chain FenAB136 and FenAB208, were generated via DNA Assembly (NEBuilder) using heavy and light chain vectors kindly provided by Mirjana Lilic. Heavy chains have a C-terminal hexa-histidine tag. Vectors encoding the Fab light chains FenAB609 and FenAB709 were generated by QuikChange site-directed mutagenesis (Agilent QuikChange Lightning site-directed mutagenesis kit) using the FenAB136 light chain as a template. The resulting expression vectors encode Fabs under the control of a CMV promoter.

### Monoclonal antibody expression and purification

For IgGs, Human Embryonic Kidney (HEK) 293F cells (ThermoFisher Scientific R79007) grown in 125 ml flasks in FreeStyle^TM^ 293 Expression Medium (ThermoFisher Scientific 12338018) were transfected at cell densities between 8.0×10^5^ to 1.0×10^6^ cells/ml with equal masses of plasmids containing the heavy chain and light chain genes using FreeStyle^TM^ MAX Reagent (ThermoFisher Scientific 16447100); both plasmids were diluted in OptiPRO^TM^ SFM (ThermoFisher Scientific 12309050). Transfected cells were incubated at 37 °C with 5% CO_2_ on a Digital Orbital Shaker (Southwest Science SBT300) operating at 120 rpm for 6 days. Supernatants were collected after 6 days and passed through a gravity chromatography column containing 1 ml of Pierce^TM^ Protein G Agarose resin (ThermoFisher Scientific 20397). The column was washed with 20 ml of wash buffer (150mM NaCl, 50mM Tris, pH 7.4) and eluted in 1 ml fractions with elution buffer (150mM NaCl, 50mM Glycine, pH 2.8) prior to neutralization with neutralization buffer (1M Tris, pH 8.0). Samples were analyzed by SDS-PAGE.

For Fabs, Fabs were produced in 293F cells by co-transfections of these cells with equal amounts of purified heavy and light chain plasmid, using 293fectin (Thermo Fisher 12347-019). Cells were incubated at 37 °C, 8% CO_2_ on an orbital shaker at 130 rpm for 5 days. Supernatants were harvested by centrifugation at 1500xg for 15 minutes, filtered, concentrated in an Amicon stirred cell and diluted (2x) in binding buffer (50 mM Sodium Phosphate, 20 mM Imidazole, 300 mM NaCl, pH8). Supernatant was then incubated with Ni-NTA agarose (Qiagen) for 1-2 hours at room temperature or overnight at 4 °C. Next, gravity flow columns were used to collect the Ni-NTA agarose and columns were washed with binding buffer. Fabs were eluted with elution buffer (50 mM Sodium Phosphate, 300 mM NaCl, 250 mM Imidazole, pH8), concentrated in centrifugal units and re-suspended in Dulbecco’s PBS. Samples were analyzed by SDS-PAGE.

### Determination of antibody affinity by biolayer interferometry and ITC

Biolayer interferometry (BLI) was performed on an Octet Red 96e system (Sartorius). Streptavidin-coated biosensors were pre-hydrated with PBS-T (phosphate buffered saline pH 7.5, 0.05% Tween-20), and baseline responses were measured in PBS-T. Biosensors were loaded with biotinylated fentanyl-hapten (where the biotin replaces the poly-gly sequence from fen-G4; biotinylated fentanyl derivative syntheses are described in (Baehr *et al*., 2020), 0.2 µg/ml for 60 sec. Association of antibody with biotinylated hapten was measured in PBS-T with 5 sample concentrations ranging from 5 nM – 200 nM for 180 sec, followed by dissociation in PBS-T for 300 sec. Isothermal Titration Calorimetry (ITC) experiments were performed using Microcal PEAQ-ITC (Malvern, UK). For all experiments PBS was used to dissolve fentanyl-citrate (Sigma, 2 mg/ml), fentanyl-hapten, and for the final purification step of Fabs. Titrations were performed at 25 °C by injecting consecutive (1-3 μl) aliquots of fentanyl or fentanyl-hapten (both 50 μM) into Fab fragment (4 –7 μM) with 120 second intervals. The binding data was corrected for the heat of dilution and fit to a one-site binding model to calculate the K_d_, and the binding parameters, N and ΔH. Binding sites were assumed to be identical.

### Determination of relative affinity by competition ELISA

Relative affinity was determined by competition ELISA. 96-well plates (Costar polystyrene high-binding plates, Corning) were coated overnight with fen-G4 conjugated to BSA, 50 ng/ml in a 50 mM carbonate buffer, pH 9.6 (Sigma). Plates were blocked with 1% gelatin for 1 hour, washed with PBS-T, and free drug was loaded onto plates as competitor. Plates were incubated with 20 ng/ml HIS-tagged Fab for 2 hours, washed, and incubated with Penta-His-biotin conjugate 1:5000 (Qiagen) overnight. Streptavidin-HRP 1:5000 (Thermo Fisher) was added, and plates were incubated for 1 hour and washed. Fab bound to plates was measured using SigmaFAST OPD substrate (Sigma), and quantitated by absorbance at 492 nm on a microplate reader (Tecan). The fentanyl-hapten structure used for this particular assay differs from the structures presented in supplementary figure 1. Specifically, we here used the “F3” hapten structure from (Robinson *et al*., 2020), with the exact same BSA-conjugation methodology described in that publication.

### Passive immunization

Five 6-8 weeks old female C57BL/6J mice per group were passively immunized by intraperitoneal injection of either 26.25 mg/kg or 3.75 mg/kg monoclonal antibody in 100 µl of PBS. One day later, mice were challenged in antinociception assays as described above.

### Crystallography of Fab-Ligand Complexes

Fabs (2-4 mg/ml) were mixed with a 2 to 3-fold molar excess of fen-G4 or fentanyl (Sigma F-013) in Dulbecco’s PBS and incubated for 1-2 hours at 4 °C. Complexes were purified by gel filtration (GE Healthcare, Superdex 200 Increase 10/300 GL column) in 10 mM HEPES, 150 mM NaCl, pH 8 and concentrated in centrifugal units to about 5 mg/ml for crystallization. All datasets were collected at the Paul Scherrer Institut (Beam line X06DA) at a wavelength of 1.0 Å, using a nitrogen stream at 100 °K. For FenAb709 iMosflm (Battye *et al*., 2011) was used to remove reflections overlapping with ice rings. All the other data sets were processed using the XDS package (Kabsch, 2010). The structure of FenAb136 was solved by molecular replacement with PHENIX (Liebschner *et al*., 2019) (PHASER module) using PDB ID 5H2B as a search model (Tatsumi *et al*., 2017). All the other structures were solved using FenAb136 as a starting model. The AUTOBUILD module of PHENIX was used for constructing the initial models. PHENIX, Coot (Emsley *et al*., 2010), and PDB-redo (Joosten *et al*., 2014) were used for model building and refinement. The fentanyl or fen-sort was built after refinement of the protein structure. In all cases the density for the ligand was clearly visible in the electron density maps.

## Supplemental Videos

**Supplemental video 1.** Fent-VAST vaccinated are protected from fentanyl effects, while control mice display the Straub Tail response.

