## Supplemental Figures for "An Epitope-Focused Trypanosome-Derived Vaccine Platform Elicits High-Affinity Antibodies and Immunity Against Fentanyl Effects"

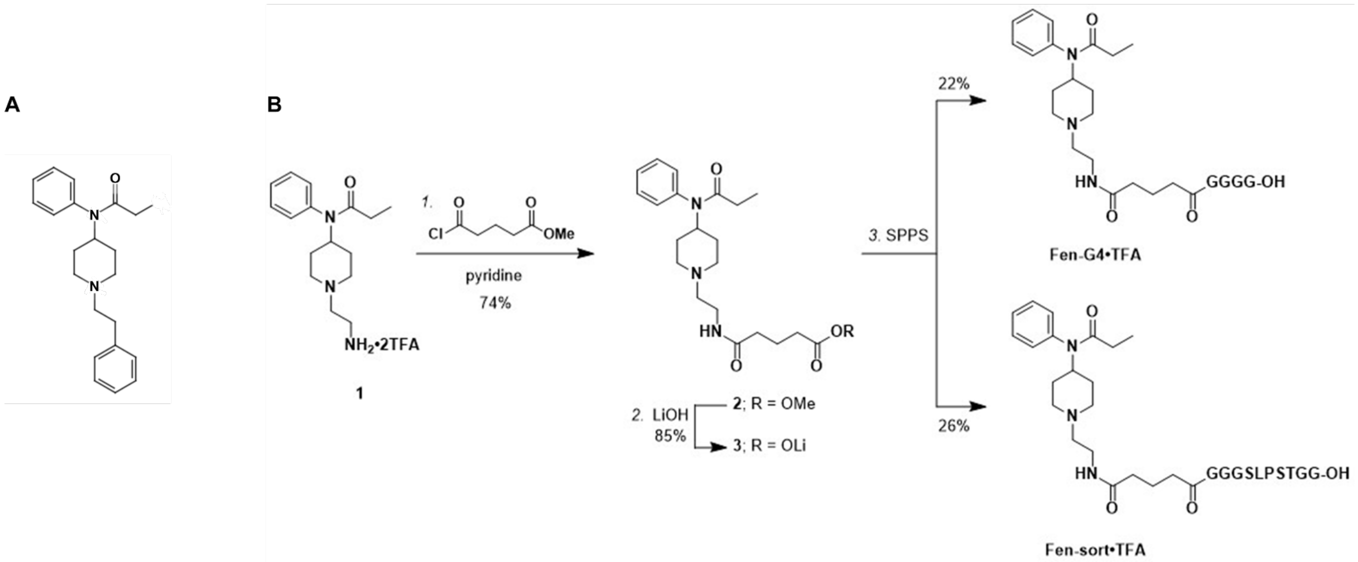


**Suppl. Fig. 1. Synthesis of fen-sort and fen-G4**

A. Chemical structure of the fentanyl molecule. Atoms are color coded as per CPK coloring. B. Synthesis of the fentanyl derivatives used in this study. Reagents and conditions: 1. glutaric acid monomethyl ester chloride (1.0 equiv), pyridine (6.0 equiv), CH2Cl2, 0 °C to rt, 16 h, 74%; 2. LiOH (3 equiv), MeOH/H2O (4:1), rt, 22 h, 85%; 3. Solid-phase peptide synthesis using an Fmoc protection strategy provided fen-G4 (22%) and fen-sort (26%) as trifluoroacetate salts (see experimental section for reaction details). Fen-sort was used to generate Fent-VASTs, while fen-G4 was used to generate fent-BSA for ELISA coating and for some of the co-crystallization studies. Both the sortaggable and poly-glycine versions of fentanyl are missing the extended aromatic ring at the bottom of the molecule. Amino acids are displayed using their single letter codes in black text. Note that this is a modified version of a previously published synthesis (Raleigh et al., 2019; Robinson et al., 2020).

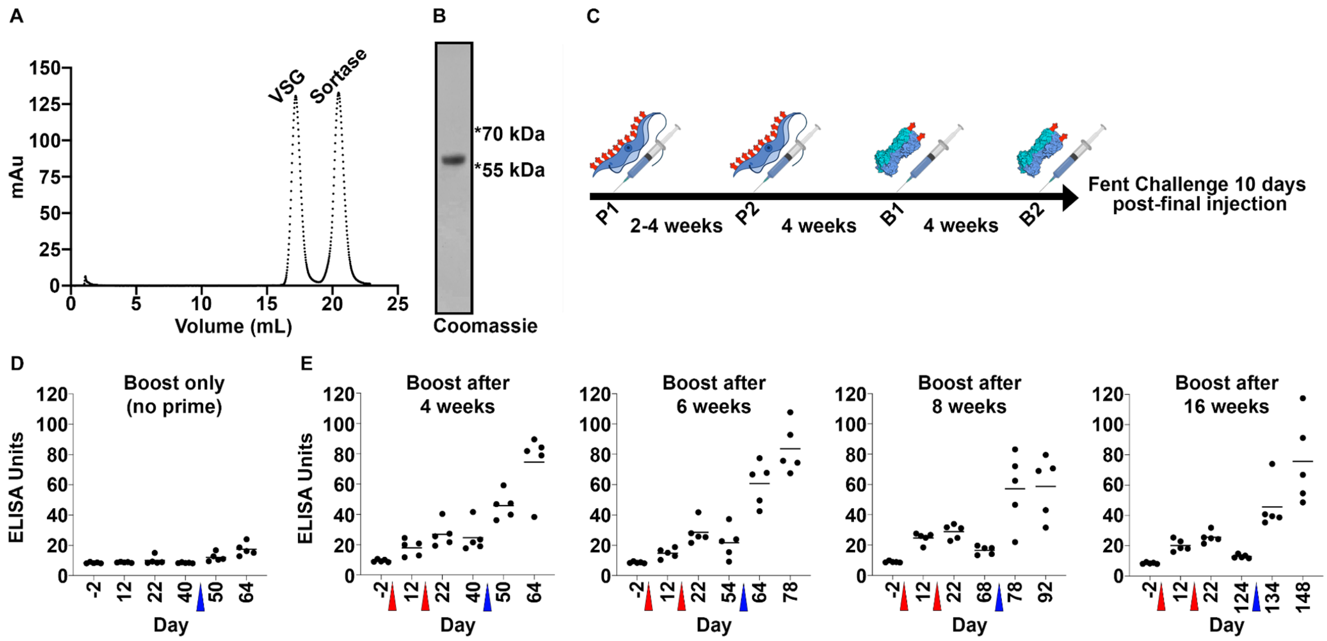


**Suppl. Fig. 2. Vaccination material, schedule, and boost-timing-dependent antibody titers**

A. Gel filtration chromatogram (from a Superose 6 increase 10/300 column; Cytiva) showing the re-purification of VSG after a sortagging reaction. The Y-axis depicts the absorbance at 280 nM, represented as milli-absorbance units (mAu). B. Coomassie-stained SDS PAGE separation of the VSG peak from (A). VSG, existing as a >55 kDa monomer, is the only observable protein in the eluted sample. C. The general vaccination schedule according to which mice represented by figures 2 and 3 were injected with Fent-VASTs, with the time intervals between each vaccination denoted between each injection day. P1 and P2; the two prime injections were composed of UV-irradiated fentanyl-coated T. brucei surface coats. B1 and B2; the two boost injections were composed of soluble Srt-VSG3 conjugated to fen-sort. Fentanyl challenge studies were always conducted 10 days post-B2. D. Anti-fentanyl antibody titers in mice immunized once (blue arrowhead) with soluble fen-sort-conjugated Srt-VSG3. E. Anti-fentanyl antibody titers in mice immunized with modified schedules as indicated above each graph. The prime injections were 2 weeks apart (at days 0 and 14), and are marked by red arrowheads. The mice were then boosted one time: 4, 6, 8, or 16 weeks after the second prime injection. The boost injections are marked by the blue arrowheads.

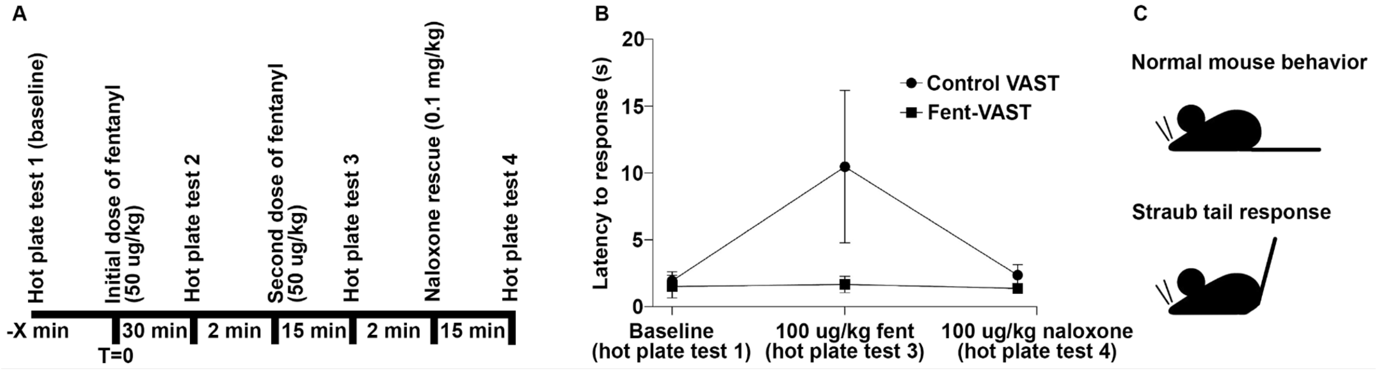


**Suppl. Fig. 3. Behavioral assays.**

A. Experimental setup of hotplate assay is shown. B. Analgesic activity was tested by using the hotplate antinociception assay as described in A. Fentanyl effect on hotplate antinociception was tested in mice immunized with control-VAST and in mice immunized with Fent-VAST. On the x-axis the baseline, the cumulative fentanyl dose received, and the naloxone dose are noted. The effect of fentanyl is shown as latency to response. Mean ± standard deviation of 5 mice per group are shown. C. Straub-Tail reaction and the position of the tail of normal mice and mice intoxicated with fentanyl is shown.

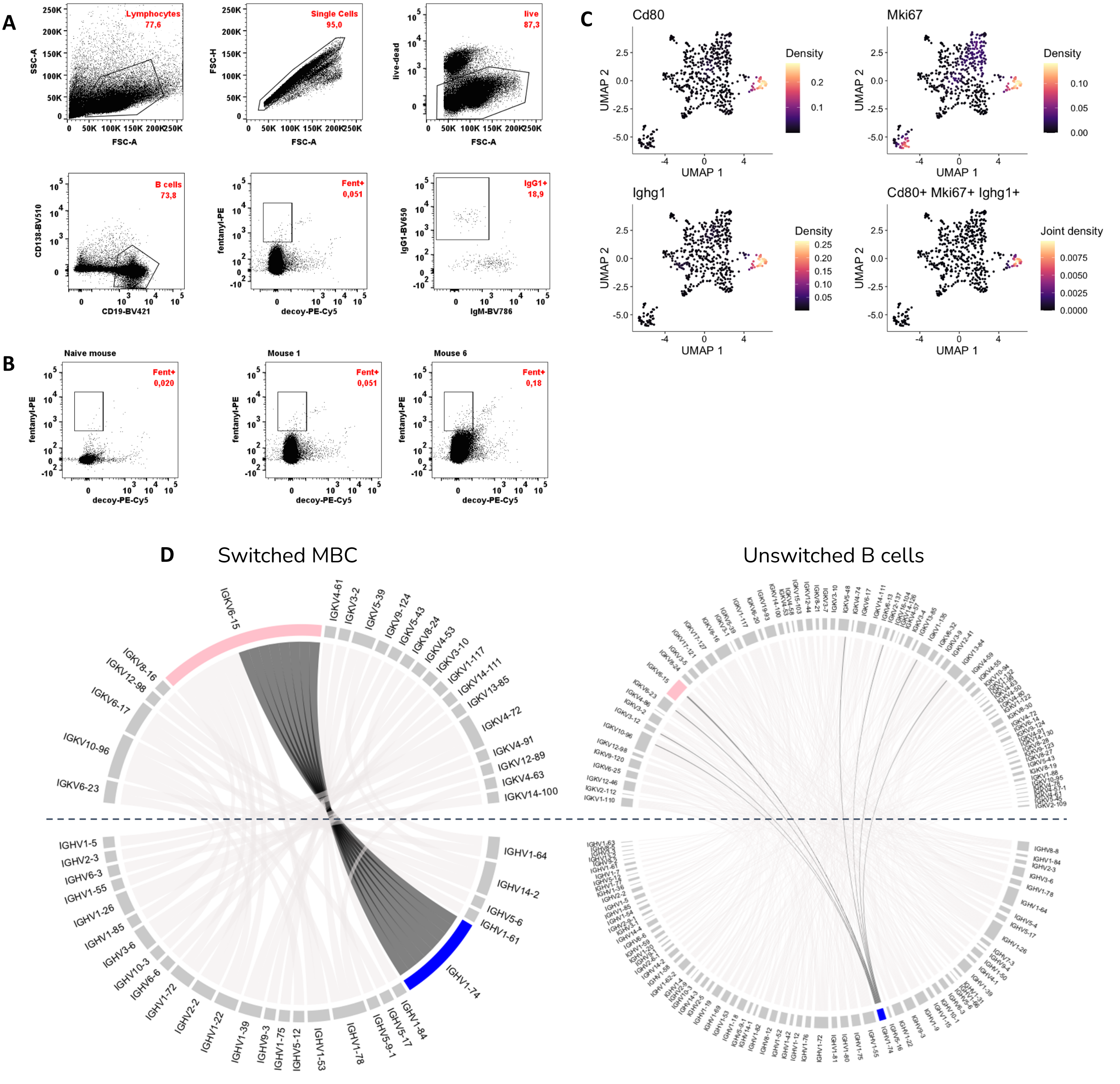


**Suppl. Fig 4. Isolation and characterization of memory B cells**

A. Gating strategy for sorting fentanyl-specific B cells from mice immunized as indicated in Fig. 1B. Splenocytes from two mice were stained with a live/dead marker, several B cell markers (CD19-BV421 and CD138-BV510), SA-BSA-PE-AF647 (decoy), and fentanyl-PE as bait. Fentanyl-PE single positive cells were sorted into 384 well plates and processed as described in the methods section. B. Sort gates for splenocytes pre-gated as shown in A for a naïve mouse and two immunized mice. The percentage of fentanyl-binding B cells in the total B cell pool is shown in red. C. UMAP visualizing the expression density of the Cd80, Mki67 and Ighg1 genes and the joint expression is shown. Each UMAP depicts the kernel density estimates of each of the selected genes, using the normalized gene expression as an input. The joint expression is calculated by multiplying the kernel density estimates from all 3 genes. Purple stands for low simultaneous expression while beige stands for high. D. Circos plots show the switched memory B cell variable region repertoire compared to non-switched B cell repertoire, for both mice combined. The expanded heavy (in blue - IGHV1-74) and light (in pink - IGKV6-15) variable regions are highlighted. In dark gray we show the pairing of the IGHV1-74 variable region of the heavy chain with the corresponding variable region of the light chains. In the switched memory B cell population (Switched MBCs) all the cells expressing a BCR with an IGHV1-74 variable region are paired with a IGKV6-15 light variable region, showing the expansion of this gene usage inside the population. This is not the case for the non-switched cells that belong to the other B cell subpopulations (right panel). There is only one cell expressing a BCR containing both an IGHV1-74 and IGKV6-15.

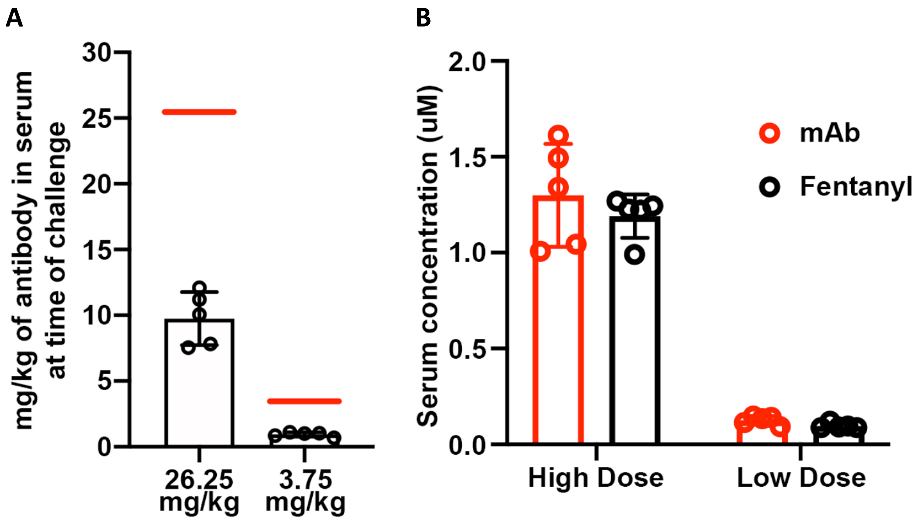


**Suppl. Fig. 5. Serum concentration of antibody before fentanyl challenge**

A. The amount of serum-circulating antibody at the time of fentanyl challenge post-passive therapy (see Fig 6E) was analyzed by western blot and plotted here. The red lines indicate the hypothetical antibody level if 100% of the injected antibody was present in the serum, highlighting the approximately 2-3-fold lower-than-maximum levels of antibody in the serum at time of challenge. B. The molar serum concentrations of anti-fentanyl IgG and fentanyl are plotted. The values suggest that approximately 95-98% of the serum-circulating antibody is able to capture fentanyl after drug injection.

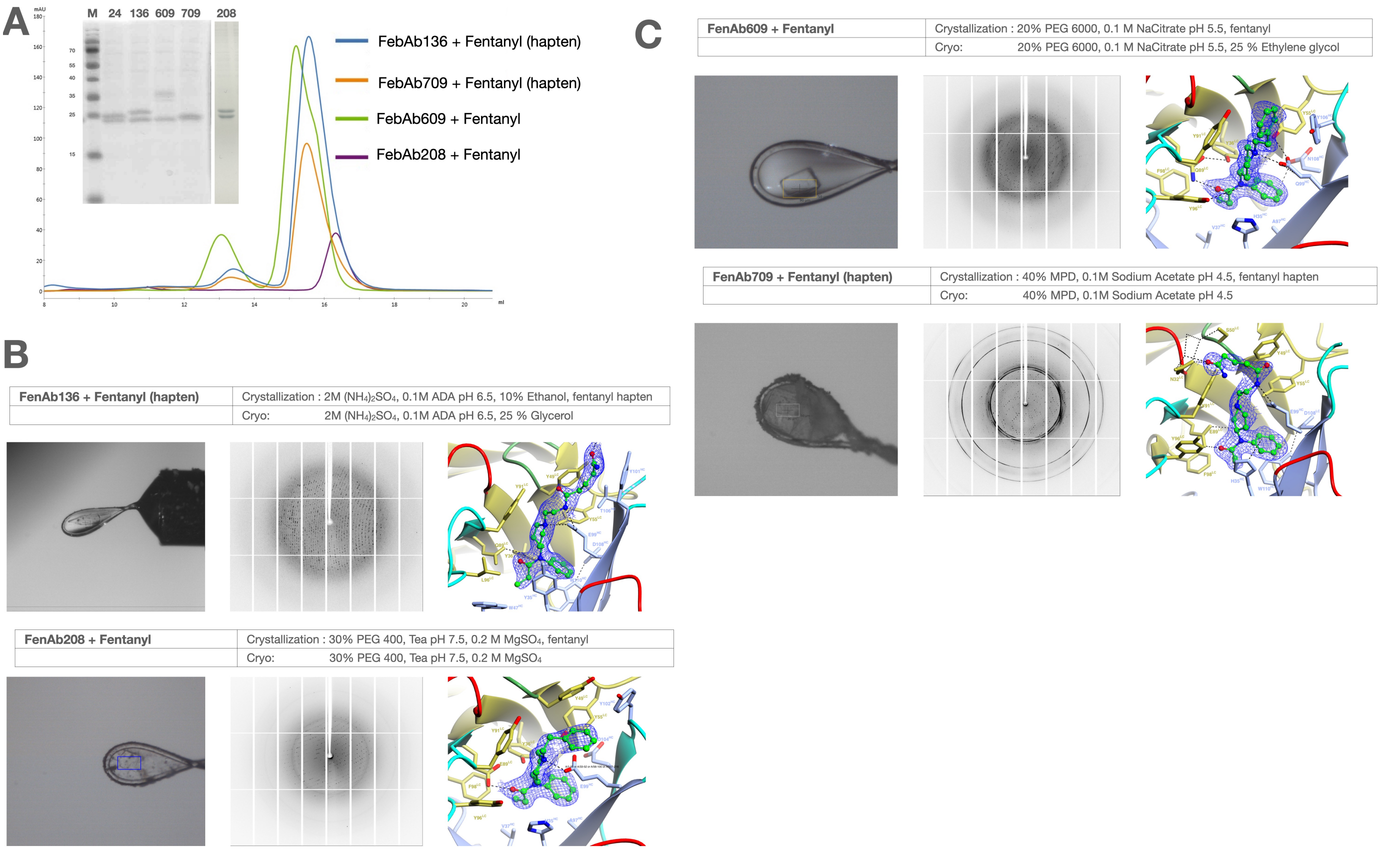


**Suppl. Fig. 6. Purification and Crystallography of Recombinant Antibody-Fentanyl Complexes**

A. Superimposed size exclusion chromatograms for final material used in crystallization experiments is shown. Elution volumes are depicted along with absorbance units at 280nm and colored for each Fab and ligand as indicated in the key. An SDS gel stained with Coomassie blue shows the contents of the peak fractions (M indicates molecular weight markers with several of those bands shown on the left side of the gel in kDa whereas the gel lanes are labeled above for the antibody number). B/C. Three panels are shown for each antibody-ligand complex with the crystallization and cooling conditions (labeled): cryo-cooled crystal mounted in loop at the synchrotron during data collection, sample diffraction, and 2Fo-Fc electron density map focused on the ligand contoured at 1 sigma from final refinements. The protein and ligand are illustrated similar to Figure 6C.

**Suppl. Table 1. Data Collection and Refinement Statistics**

| | FenAb136 | FenAb208 | FenAb609 | FenAb709 |
|---|---|---|---|---|
| **Wavelength (Å)** | 1.0 | 1.0 | 1.0 | 1.0 |
| **Resolution range (Å)** | 54.52 - 2.07 (2.14 - 2.07) | 48.49 - 1.92 (1.99 - 1.92) | 45.19 - 1.7 (1.76 - 1.70) | 55.32 - 2.32 (2.40 - 2.32) |
| **Space group** | C 1 2 1 | P 41 | P 21 21 21 | C 2 2 21 |
| **Unit cell Dimensions (Å)** | 182.98 105.84 172.77 90 112.24 90 | 153.34 153.34 45.99 90 90 90 | 77.27 111.41 114.81 90 90 90 | 75.19 138.69 101.07 90 90 90 |
| **Total reflections** | 361101 (35110) | 414373 (39287) | 727900 (70440) | 243094 (25540) |
| **Unique reflections** | 181970 (17881) | 82358 (8154) | 109391 (10846) | 22297 (2274) |
| **Multiplicity** | 2.0 (2.0) | 5.0 (4.8) | 6.7 (6.5) | 10.9 (11.2) |
| **Completeness (%)** | 97.51 (96.49) | 99.85 (99.88) | 99.89 (99.71) | 94.77 (100.00) |
| **Mean I/sigma(I)** | 7.29 (1.20) | 11.69 (1.05) | 11.31 (0.95) | 16.19 (7.30) |
| **Wilson B-factor (Å$^2$)** | 35.64 | 33.63 | 25.77 | 25.34 |
| **R-merge** | 0.0637 (0.5982) | 0.08342 (1.312) | 0.09267 (1.943) | 0.09728 (0.2856) |
| **R-meas** | 0.09008 (0.8459) | 0.09322 (1.475) | 0.1006 (2.111) | 0.1021 (0.2991) |
| **R-pim** | 0.0637 (0.5982) | 0.04106 (0.6656) | 0.03887 (0.819) | 0.03055 (0.08823) |
| **CC1/2** | 0.987 (0.584) | 0.999 (0.577) | 0.999 (0.608) | 0.998 (0.978) |
| **CC*** | 0.997 (0.859) | 1 (0.856) | 1 (0.87) | 0.999 (0.995) |
| **Reflections used in refinement** | 180911 (17746) | 82332 (8148) | 109323 (10826) | 22035 (2274) |
| **Reflections used for R-free** | 1986 (185) | 4117 (408) | 5470 (542) | 1110 (149) |
| **R-work** | 0.2424 (0.3486) | 0.1960 (0.3754) | 0.2015 (0.4464) | 0.2096 (0.2261) |
| **R-free** | 0.2816 (0.3953) | 0.2299 (0.3960) | 0.2302 (0.4786) | 0.2446 (0.2859) |
| **CC(work)** | 0.921 (0.540) | 0.966 (0.788) | 0.962 (0.810) | 0.932 (0.896) |
| **CC(free)** | 0.904 (0.536) | 0.960 (0.720) | 0.942 (0.808) | 0.910 (0.784) |
| **Number of non-hydrogen atoms** | 21106 | 7011 | 7371 | 3469 |
| **macromolecules** | 19606 | 6550 | 6822 | 3180 |
| **ligands** | 172 | 50 | 50 | 28 |
| **solvent** | 1328 | 411 | 499 | 261 |
| **Protein residues** | 2558 | 852 | 855 | 415 |
| **RMS(bonds) (Å)** | 0.002 | 0.011 | 0.010 | 0.004 |
| **RMS(angles) (Å)** | 0.52 | 1.14 | 1.06 | 0.72 |
| **Ramachandran favored (%)** | 97.94 | 98.21 | 98.46 | 97.27 |
| **Ramachandran allowed (%)** | 2.06 | 1.67 | 1.54 | 2.73 |
| **Ramachandran outliers (%)** | 0.00 | 0.12 | 0.00 | 0.00 |
| **Rotamer outliers (%)** | 1.22 | 0.93 | 0.89 | 3.57 |
| **Clashscore** | 2.86 | 3.09 | 1.71 | 3.68 |
| **Average B-factor (Å$^2$)** | 45.82 | 43.26 | 37.22 | 32.42 |
| **macromolecules** | 45.85 | 43.22 | 37.20 | 32.69 |
| **ligands** | 46.12 | 41.48 | 30.41 | 27.19 |
| **solvent** | 45.40 | 44.17 | 38.13 | 29.67 |
| **Number of TLS groups** | 50 | 18 | 14 | 8 |
| Statistics for the highest-resolution shell are shown in parentheses. | | | | |
